# Impaired bidirectional communication between interneurons and oligodendrocyte precursor cells affects cognitive behavior

**DOI:** 10.1101/2021.05.04.442422

**Authors:** Lipao Fang, Na Zhao, Laura C. Caudal, Renping Zhao, Ching-Hsin Lin, Hsin-Fang Chang, Nadine Hainz, Carola Meier, Bernhard Bettler, Wenhui Huang, Anja Scheller, Frank Kirchhoff, Xianshu Bai

## Abstract

Cortical neural circuits are complex but very precise networks of balanced excitation and inhibition (E/I). Yet, the molecular and cellular mechanisms that form the E/I balance are just beginning to emerge. Here, using conditional GABA_B_ receptor-deficient mice we identified a GABA/TNF-related cytokine (TNFSF12)-mediated bidirectional communication pathway between Parvalbumin-positive (PV^+^) fast spiking interneurons and oligodendrocyte precursor cells (OPCs) that determines the density and function of interneurons in the developing medial prefrontal cortex (mPFC). Interruption of the GABAergic signaling to OPCs resulted in reduced myelination and hypoactivity of interneurons, strong changes of cortical network activities and impaired cognitive behavior. In conclusion, glial transmitter receptors are pivotal elements in finetuning distinct brain functions.

## Introduction

The inhibition of cortical network activity is performed by interneurons that release the inhibitory transmitter γ-aminobutyric acid (GABA) which acts on ionotropic GABA_A_ and metabotropic GABA_B_ receptors (GABA_B_R) ^1, 2^. Alterations of interneuron cell density as well as concomitant changes of firing activity are often observed in several neuropsychiatric conditions ^3–6^. In the mouse cortex, higher frequencies of spontaneous inhibitory postsynaptic currents (sIPSCs) were observed after increasing the interneuron density by transplantation of precursor cells ^7^. Similarly, blocking of interneuron apoptosis caused a decrease of the excitation/inhibition (E/I) ratio ^6, 8^. Interneuron density is a pivotal determinant of correct inhibitory circuits in the central nervous system (CNS). During development, a surplus of interneurons is generated that populates the cortical plate ^9^. Subsequently, about 40 % of cortical inhibitory neurons are eliminated by programmed cell death during the first two postnatal weeks ^7^. One hypothesis links the apoptosis of supernumerary neurons with the onset of first neuron-neuron connections ^10^. Only the neurons that receive sufficient neurotrophic signals from their target cells will survive ^10, 11^.

Oligodendrocyte precursor cells (OPCs) form synapses with interneurons as early as postnatal day (p) 4-5 ^12^. Concomitantly, the incidence of interneuron apoptosis increases drastically after p5 and reaches its peak at p7 ^7^. Right after, the interneuron-OPC connectivity accelerates till p10, which drives an immediate oligodendrocyte (OL) boom and subsequent interneuron myelination via OPC-GABA_A_ and GABA_B_ receptors ^13–15^. Despite relatively short axons, myelination of cortical interneurons (mainly parvalbumin (PV)^+^ fast-spiking interneurons, ^16^) contributes significantly to the fine-tuning of local activity in the medial prefrontal cortex (mPFC) and cortex, ensuring proper performance of behavior ^4, 17^. However, how the early communication between interneuron and OPC affect inhibitory network activity in the mPFC still remains unknown.

To assess the function of OPC GABA_B_Rs for mPFC inhibition, we conditionally deleted the GABA_B_ receptor subunit *gabbr1* selectively in OPCs at the end of the first postnatal week. By focusing on the medial prefrontal cortex (mPFC), we found that OPCs shape the inhibitory network by GABA_B_ receptor and the cytokine tumor necrosis factor-like weak inducer of apoptosis (TWEAK) signaling pathways, thereby finetuning interneuron cell death and survival as well as its myelination onset. The functional and morphological changes of PV^+^ interneurons observed in the adult mPFC of mutant mice generate a reduction of the inhibitory tone, which finally leads to impaired cognition and social behavior.

## Results

### GABA_B_Rs of OPCs are required for oligodendrogenesis in the mPFC

To assess the function of GABA_B_Rs for OPC-interneuron communication, we generated Tg*(NG2-CreER^T2^):GABA_B1_R^fl/fl^* mice (Fig. 1a) to conditionally knockout (cKO) functional GABA_B_R from OPCs and their progeny. We induced the cKO at postnatal day 7 and 8 (p7/8) before the onset oligodendrocyte formation at p10-11 ^13^. When analyzed at the age of 9 weeks (w), about 80 % of OPCs (platelet-derived growth factor receptor α^+^ (PDGFRα^+^, Pα+))were found recombined (Pα^+^tdT^+^/Pα^+^) in the medial prefrontal cortex (mPFC), based on tdTomato (tdT) gene expression (Fig. 1b, c).

**Figure 1.**
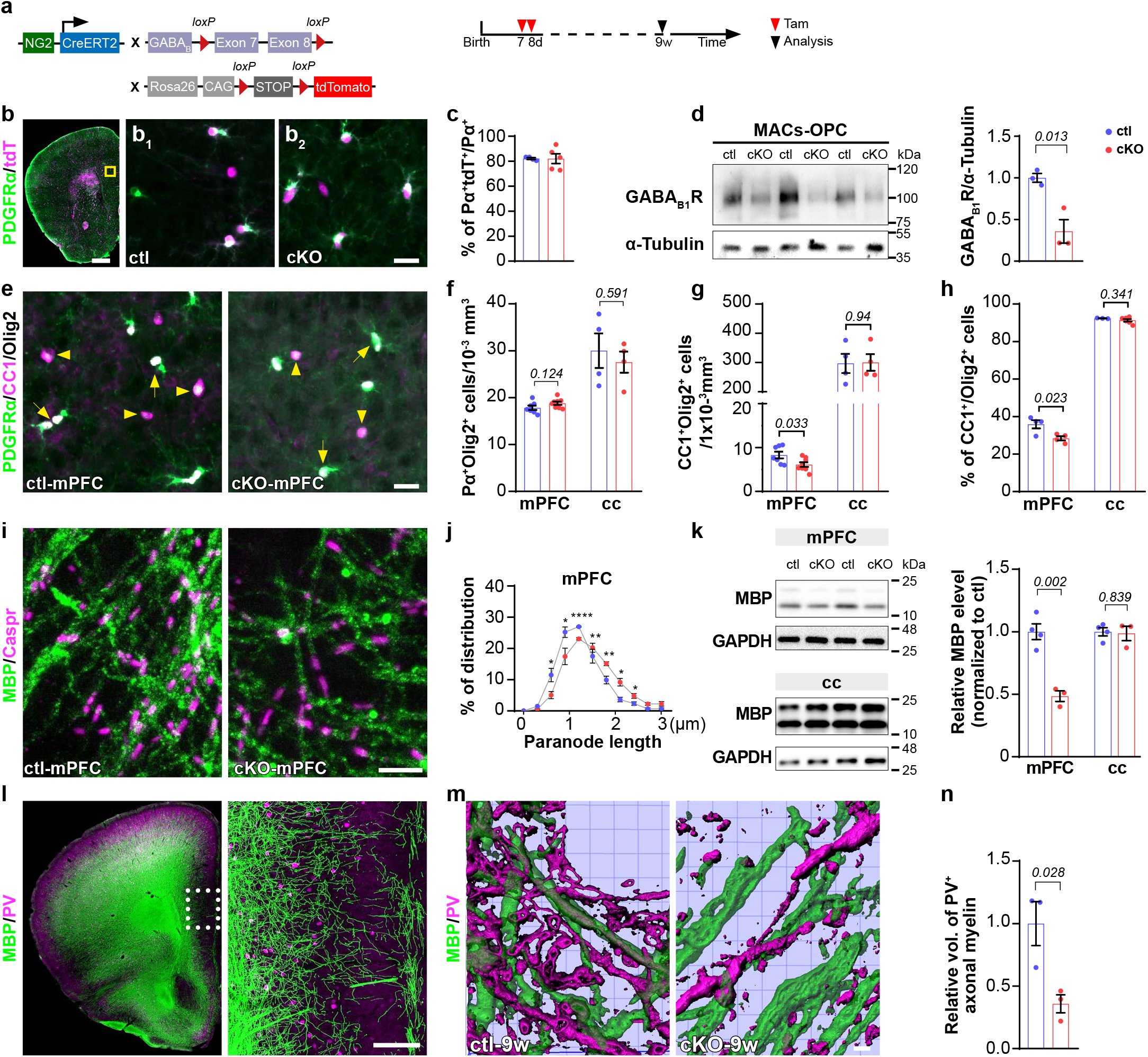
Ablation of GABA_B_Rs in OPCs attenuates oligodendrocyte differentiation and myelination of interneurons in the medial prefrontal cortex. ***a***, Genetic strategy and experimental schedule. ***b***, Prefrontal cortex in coronal section stained with PDGFRα. tdTomato^+^ (tdT^+^) cells indicate recombined cells. ***b_1_***, ***b_2_***, magnified images of ctl and cKO medial prefrontal cortex (mPFC, highlighted in *b*). ***c***, Recombination efficiency of OPCs in ctl and cKO mPFC. (*ctl=4, cKO=5, unpaired t-test*). ***d***, Western blot analysis of GABA_B_R subunit 1 in MACs sorted OPCs from cortex. (*ctl=3, cKO=3, unpaired t-test*). ***e***, Exemplary images of Olig2, PDGFRα and CC1 immunostaining in adult ctl and cKO mPFC. ***f, g***, Quantification of OPC and oligodendrocyte cell densities in mPFC and corpus callosum (cc). (mPFC: *ctl=7, cKO=8, cc: ctl=4, cKO=4, multiple t-tests)*. ***h***, Quantification of the oligodendrocyte proportion among the total lineage (mPFC: *ctl=4, cKO=4;* cc: *ctl=3, cKO=6, multiple t-tests)*. ***i***, Immunostaining of Caspr and MBP in ctl and cKO mPFC. ***j***, Quantitative analysis of paranode length in mPFC (*ctl=4, cKO=4, multiple t-tests*). ***k***, Western blot analysis of MBP expression in mPFC and cc. (*ctl=4, cKO=3, multiple t-tests*). ***l***, Overview of coronal brain slices immunostained with parvalbumin (PV) and MBP shows abundant PV^+^ neurons are myelinated in mPFC. ***m***, 3D reconstruction of myelinated PV^+^ axons in ctl and cKO mPFC using Imaris software. ***n***, Relative volume of MBP^+^ myelin sheaths wrapping PV^+^ axons (*normalized to ctl). (ctl=3, cKO=3, unpaired t-test*). Scale bars in b=1 mm, b_2_, e=10 µm, i, m=5 µm, l=200 µm.

To confirm and quantify the deletion of GABA_B_R in OPCs, we performed Western blot analysis with OPCs purified by magnetic-activated cell sorting (MACs-OPCs). In cKO OPCs, GABA_B_R expression was reduced by about 64 % compared with controls (ctl) (Fig. 1d). Considering the substantial purity of MACS-sorted OPCs (85 %, data not shown) and recombination efficiency (80 %), we concluded that a vast majority of GABA_B_Rs had been ablated from OPCs.

Next, we focused on the oligodendrocyte lineage and evaluated the contribution of GABA_B_Rs for OPC differentiation in the mPFC by immunostaining of PDGFRα and adenomatous polyposis coli clone 1 (CC1) (Fig. 1e), established markers for OPCs and oligodendrocytes, respectively. In the cKO mice, the density of OPCs (PDGFRα^+^Olig2^+^) did not change (Fig. 1f), while that of mature oligodendrocytes (CC1^+^Olig2^+^) was strongly reduced to 74 % (Fig. 1g), indicative of decreased OPC differentiation to oligodendrocytes (Fig. 1h). Notably, in the corpus callosum the densities of oligodendrocytes and OPCs, as well as their relative proportions did not differ between ctl and cKO mice (Fig.1 f-h, Extended Data Fig. 1). These data suggest that OPC-GABA_B_Rs are required for OPC differentiation in the grey matter mPFC, but not in the white matter corpus callosum.

We further analyzed structural aspects of myelination which is important for axonal conductivity, i.e. the lengths of nodes (the gaps between paired contactin-associated protein-positive (Caspr^+^) segments) and paranodes (single Caspr^+^ segments) by immunostaining (Fig. 1i). In cKO mPFC, the paranodal length was increased (Fig. 1j) while node length and density remained stable (Extended Data Fig. 2). In parallel, we observed a reduction of myelin basic protein (MBP) expression at the mRNA (Extended Data Fig. 2) and protein levels (Fig. 1k) in cKO mPFC, while the other myelin protein, proteolipid protein (PLP) mRNA levels were not changed (Extended Data Fig. 2). In the corpus callosum, however, myelination was unperturbed as indicated by unchanged paranode length (Extended Data Fig. 2), node density (Extended Data Fig. 2) and MBP expression (Fig. 1k). Therefore, our data show that GABA_B_Rs of OPCs promote differentiation and myelination in the mPFC, but do not affect the corpus callosum.

Since OPCs continuously differentiate into oligodendrocytes ^18–20^, the cKO mice will also generate oligodendrocytes lacking GABA_B_Rs. To selectively target only oligodendrocytes, we used Tg*(PLP-Cre^ERT2^):GABA_B1_R^fl/fl^* mice (Extended Data Fig. 3). The gene deletion was induced by tamoxifen injections at p7/8 or at 4w. At the age of 9w, we did not observe differences in the OPC and oligodendrocyte population between ctl and cKO mice (Extended Data Fig. 3), neither in the PFC nor in the corpus callosum (Extended Data Fig. 3).

In summary, our data strongly suggest that GABA_B_R is specifically required for early OPC differentiation and subsequent myelination in the mPFC, but not in the corpus callosum.

### GABA_B_Rs of OPCs promote the myelination of interneurons in the mPFC

Since callosal axons are primarily excitatory ^21^ and axons of inhibitory interneurons are largely restricted to cortical areas of the same hemisphere (including mPFC, Fig.1l) (Extended Data Fig. 4), we hypothesized that the observed myelin deficits were restricted to interneurons. Required by their fast-spiking activity, parvalbumin (PV)^+^ interneurons are the most abundantly myelinated interneurons ^16, 22–24^ of the cortex. These neurons are most prominent at the layer 4 and 5 with majority of their axons projecting to the layer 2/3 and 5 ^25^ (Fig. 1l, Extended Data Fig. 4). To evaluate their myelination, we performed PV and MBP double immunostaining (Fig. 1l-n) and analyzed the volume of myelin sheaths, by volume rendering of MBP immunolabel covering PV^+^ axons at the cortical layers 2/3 of mPFC. Indeed, the volume of myelin sheaths wrapping PV neurons was reduced by 60 % in the cKO mPFC (Fig. 1n). However, the myelination of excitatory neurons (in the corpus callosum) was not affected as the g-ratio of myelinated axons and conduction velocity of compound action potential did not differ between ctl and cKO mice (Extended Data Fig. 2). Thus, we concluded that GABA_B_R of OPCs are preferentially involved in the myelination of inhibitory rather than excitatory neurons.

### Postnatal deletion of GABA_B_Rs in OPCs causes enhanced interneuron density in the adult, but with reduced GABA release

While investigating the interneuronal myelination, we recognized that early ablation of GABA_B_R in OPCs (at p7/8, Fig. 2a) was associated with a 30 % increase of interneuron density (10.4 vs 13.3 for ctl and cKO respectively) in the cortex as revealed by immunostaining for PV (Fig. 2b). However, in the mPFC this increase did not result in a concomitant increase of inhibitory input (Fig. 2c, d). In contrast, the frequency of spontaneous inhibitory postsynaptic currents (sIPSCs) of pyramidal neurons in the layer V of mPFC, where interneurons are abundantly located, was reduced by 40 %, while the amplitude of sIPSCs remained unaffected in the mutant mPFC as well as the firing rate and amplitude of excitatory postsynaptic currents (EPSCs).

**Figure 2.**
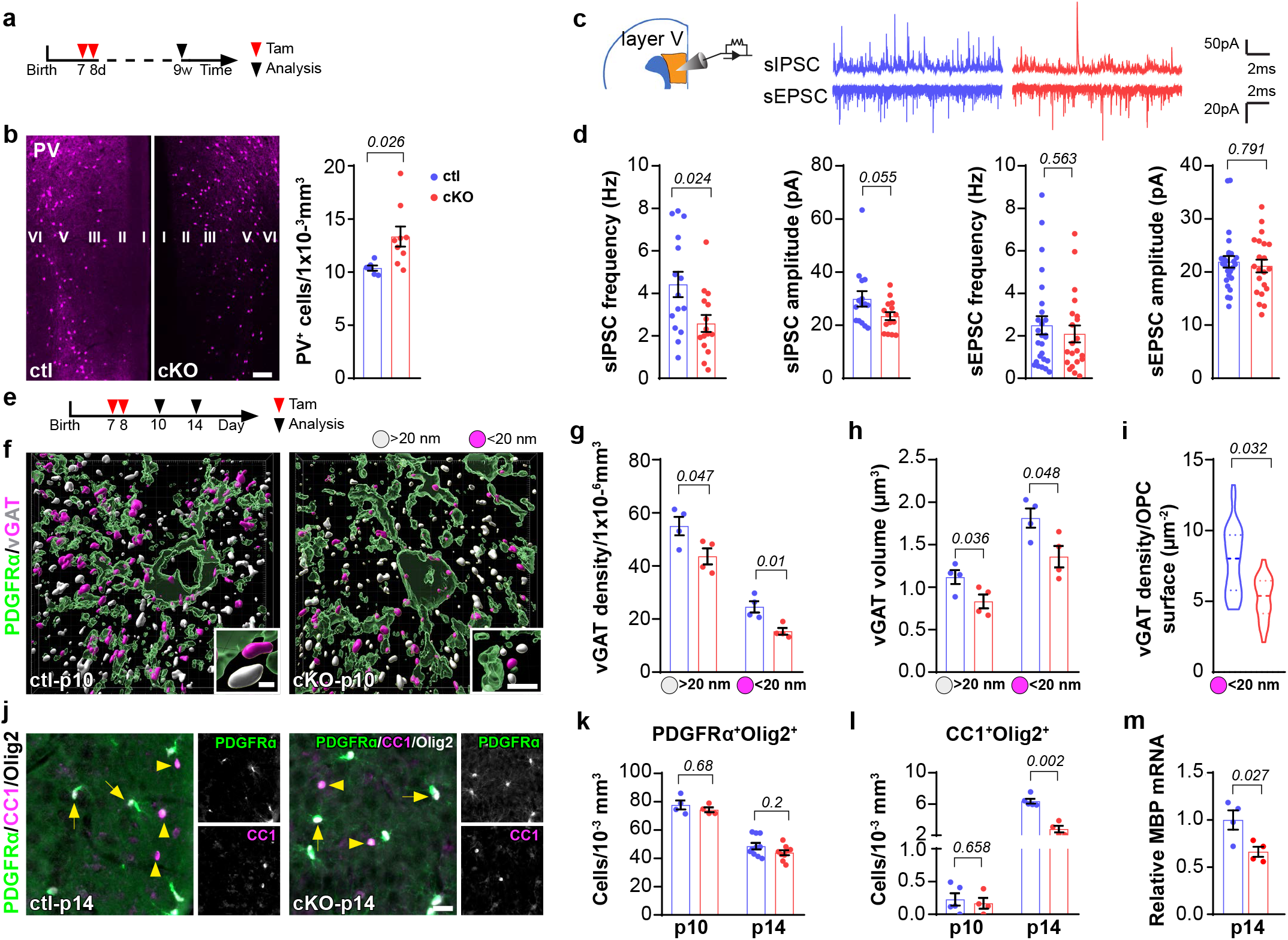
In OPC-GABA_B_R cKO mice impaired control of interneuronal density and firing rate reduces oligodendrocyte differentiation and myelin gene expression at the onset of myelination. ***a,*** Experimental schedule for ***b-d***. ***b***, Immonostaining (left) and quantification (right) of PV^+^ neurons in the mPFC at the age of 9 weeks (*ctl=5, cK=9, unpaired t-test*). ***c***, Exemplary traces of sIPSCs and sEPSCs recorded at the layer V of mPFC. ***d***, Quantification of frequency and amplitude of sIPSC and sEPSC *(sIPSC: ctl=15 cells from 4 mice, cKO=15 cells from 5 mice; sEPSC: ctl=26 cells from 4 mice, cKO=21 cells from 5 mice, Mann Whitney test). **e**,* Experimental schedule for ***f-m***. ***f***, 3D reconstruction of OPCs and vGAT by PDGFRα and vGAT immunostaining in ctl and cKO mPFC at p10 using Imaris software. vGAT was classified into two subgroups: magenta vGATs have < 20nm distance from OPC surface while grey vGATs have > 20nm. ***g, h***, Density (***g***) and volume (***h***) of vGAT in ctl and cKO mPFC. (*ctl=4, cKO=4; multiple t-tests*). ***i***, Quantification of synaptic vGAT (< 20nm) density per µm^2^ of OPC surface. *(ctl=4, cKO=4, unpaired t-test*). ***j***, Immunostaining of OPCs and oligodendrocytes with PDGFRα, Olig2 and CC1 in p14 mPFC. ***k, l***, Density of OPCs (***k***) and oligodendrocytes (***l***) in ctl and cKO mPFC at p10 and p14. (*p10 and p14: ctl=5, cKO=4; multiple t-tests*). ***m***, Quantitative analysis of MBP mRNA level in ctl and cKO mPFC at p14. (*ctl=4, cKO=4, unpaired t-test)*. Scale bars in **b**=50 µm, **f**=5 µm, **j**=10 µm.

Since neuronal activity can regulate myelination ^22, 26^, we asked whether the reduced inhibitory firing rate contributed to the observed molecular and structural changes of the myelin at interneuronal axons in the mutant mPFC. For that purpose, we further analyzed the cKO mPFC of p10 mice (Fig. 2e) at the onset of myelination and when interneuron-OPC communication peaks ^13^. After induction of the cKO at p7/8, we assessed the density of the vesicular GABA transporter (vGAT) close to OPC surfaces as a quantitative readout of inhibitory presynaptic terminals by combined vGAT and PDGFRα immunostaining ^27^ (Fig. 2f-i, Extended Data Fig. 5). We classified the vGAT puncta into two subgroups based on the distance between vGAT staining and OPC surface: ‘<20 nm’ as vesicles targeting on postsynaptic OPCs (magenta), and ‘>20 nm’ as vesicles targeting on other cells (grey), assuming a width of about 20 nm for the synaptic cleft ^28^. And indeed, already at p10, i.e. only 3 days after the first tamoxifen injection, the density and the volume of vGAT immunopuncta were reduced were reduced to 75 % and 80 % in the cKO mPFC (Fig. 2f-i, Extended Data Fig. 5), including a smaller volume of vGAT (Fig. 2h) and less puncta (Fig. 2i) at the OPC surface in cKO mPFC, which can explain the attenuated firing rate of the interneurons in the adult. These results indicate that interneurons exhibit lower activity and transmit less GABAergic signals to OPCs in the cKO mPFC.

To test whether the reduced inhibitory tone affecting OPC differentiation specifically correlates with the onset of myelination, we compared the OPC and oligodendrocyte populations at p10 and p14 (Fig. 2j-m, Extended Data Fig. 6). At p10, the densities of OPC and oligodendrocyte populations were still comparable in the cKO mPFC to control mPFC, although the interneurons already expressed less vGAT (Fig. 2f-h). However, at p14, the differentiation and cell numbers of oligodendrocytes dropped strongly in the mutant mPFC (Fig. 2l, Extended Data Fig. 6), as did the MBP expression (Fig. 2m). Again, in the corpus callosum, OPC and oligodendrocyte population were not affected by the deletion of GABA_B_Rs (Extended Data Fig. 1).

When we induced GABA_B_R deletion in the young adult mice at the age of 4 weeks, both in the mPFC and corpus callosum, the density of oligodendrocyte as well as the MBP expression were not affected in the cKO mice (Extended Data Fig. 7).

In summary, our data strongly suggest that during early development of the mPFC deletion of GABA_B_R in OPCs affects the inhibitory tone of interneurons, feeding back onto OPC differentiation and interneuron myelination.

### OPCs induce interneuron apoptosis via GABA_B_R-TWEAK signaling

As we had seen a surplus of interneurons in the mutant mPFC which could explain the abnormal neuronal activity ^8, 29^, we tested whether the interneuron numbers would be accurately controlled by programmed cell death early during development. After induction of GABA_B_R in OPCs at p7/8, the extent of interneuron apoptosis was determined at p10 and p14 by immunostaining for PV and cleaved caspase-3 (CC-3, Extended Data Fig. 8), a well-established marker for apoptosis. As expected by the earlier experiments, we found 50 % reduction of PV^+^/CC-3^+^ interneurons at p10 suggestive of strongly reduced apoptosis of PV^+^ interneurons in the mutant mPFC (Extended Data Fig. 8). Obviously, OPCs receive GABAergic input through GABA_B_R and send back pro-apoptotic signals to interneurons.

To confirm the selective cell death of interneurons, we induced the cKO as early as p1 and p2 and compared the apoptosis of inhibitory neurons with that of excitatory ones at p5 and p7 (Fig. 3a). Interneuron apoptosis persists from p1 till p15 with a peak at p7 ^7^, while the time window for excitatory neuron apoptosis is rather narrow, between p2-5 ^30^. Apoptosis of total interneurons (GAD67^+^CC-3^+^) as well as PV interneurons (PV^+^CC-3^+^ left panel in Fig. 3b) was significantly reduced in cKO mPFC at p5 and p7 (Fig. 3c), while that of the excitatory neurons (CTIP^+^CC-3^+^, right panel in Fig. 3b; TBR1^+^CC-3^+^) did not change at both time points (Fig. 3c, Extended Data Fig. 9). Therefore, we concluded that OPCs send pro-apoptotic signals back to interneurons at the early postnatal weeks.

**Figure 3.**
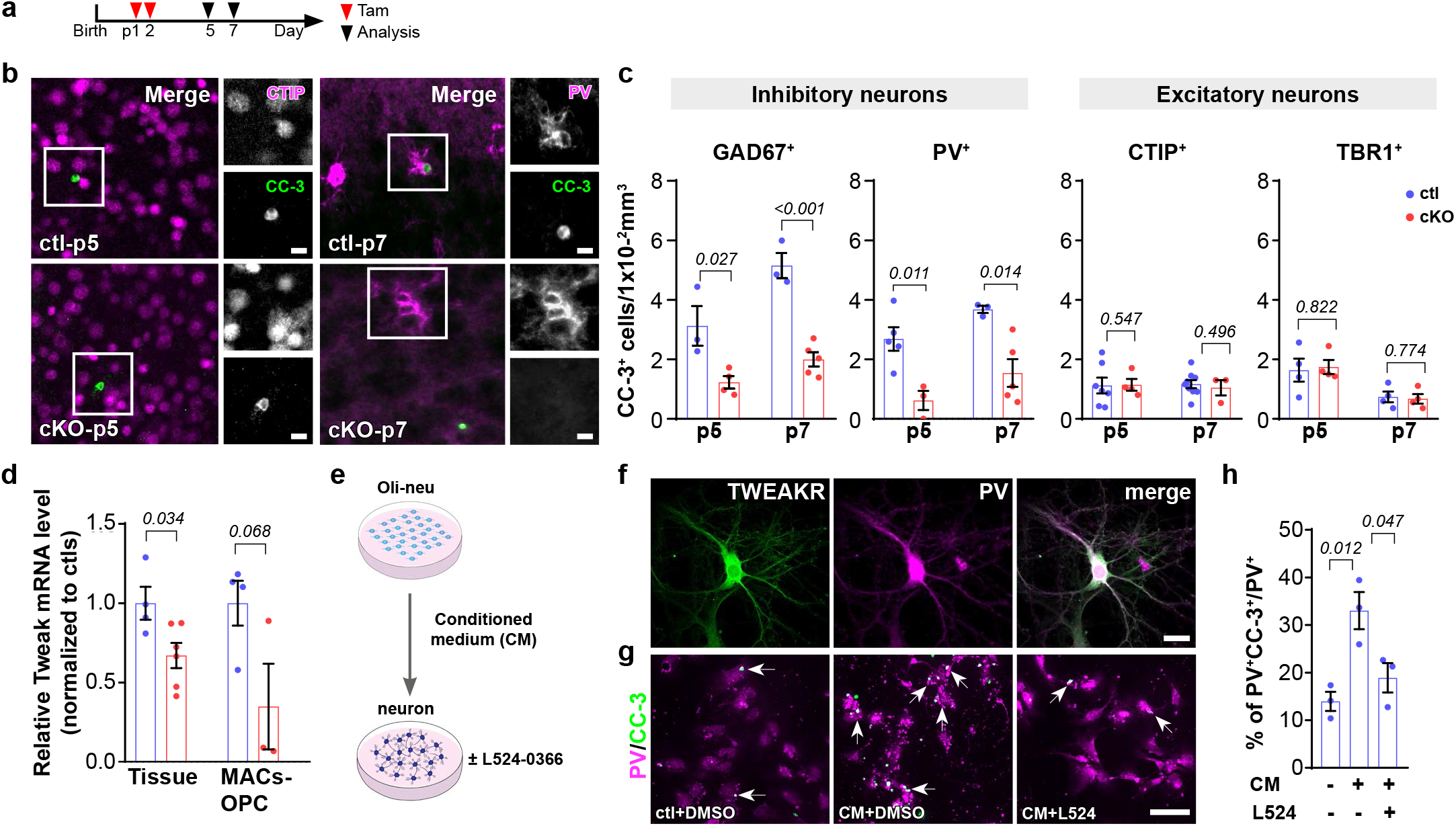
During early CNS development the apoptosis of supernumerous interneurons is mitigated upon impaired release of TWEAK from GABA_B_R-deficient OPC. ***a***, Scheme of experimental plan. ***b***, Representative images of excitatory (CTIP) and inhibitory (PV^+^) neurons co-stained with apoptotic marker CC-3 in p5 and p7 mPFC. ***c***, Quantification of apoptotic neurons (CC-3^+^) co-expressing interneuron marker GAD67 and PV or excitatory neuronal marker CTIP and TBR1. *(GAD67: p5-ctl=3, p5-cKO=4; p7-ctl=3, p7-cKO=5) (PV: p5-ctl=5, p5-cKO=3; p7-ctl=3, p7-cKO=5) (CTIP: p5-ctl=7, p5-cKO=4; p7-ctl=9, p7-cKO=3) (TBR1: p5-ctl=4, p5-cKO=4; p7-ctl=4, p7-cKO=4) (multiple t-tests)*. ***d***, Relative mRNA level of TWEAK in the mPFC and MACs-OPCs from ctl and cKO mice *(tissue-ctl=4, tissue-cKO=6; MACs-ctl=4, MACs-cKO=3; multiple t-test). **e**,* Experimental design for *in vitro* studies. ***f***, Immunolabeling of TWEAK receptor on 14 DIV PV^+^ interneurons. ***g***, Immunostaining of apoptotic primary PV^+^ neurons treated with conditional medium of Oli-neu cells co-treated with or without TWEAKR antagonist L524-0366 (20 µM). ***h***, Percentage of apoptotic PV^+^ neurons among total PV^+^ neurons after the treatments *(n=3 independent experiments, one-way ANOVA, multiple comparison)*. Scale bars in **b**=5 µm, **f**=20 µm, **g**=50 µm.

To molecularly identify the critical apoptotic factor released by OPCs, we first tested the expression levels of several peptides which had been associated with cell survival or cell death in ctl and cKO mPFC (Extended Data Fig. 10). Gene deletion was induced at p1/2, and mPFC was collected at p7, the peak of interneuron apoptosis (Extended Data Fig. 10). We found only the expression of the cytokine tumor necrosis factor-like weak inducer of apoptosis (TWEAK, also known as Apo3l or TNF superfamily member 12, TNFSF12) to be diminished about 30 % in cKO mPFC (Fig. 3d, Extended Data Fig. 10). TWEAK induces an apoptotic cell death after binding to its cognate TWEAK receptor (TWEAKR; TNF Receptor Superfamily Member 12A, TNFRSF12A; also known as FN14) ^31^. Subsequently, we tested MACS-sorted OPCs (collected from cKO and ctl cortices at p7) (Fig. 3d) as well as Oli-neu cells (an OPC cell line) treated with GABA_B_R antagonist CGP 55845 for TWEAK. Also here, we observed TWEAK downregulation (Extended Data Fig. 10). Three independent experiments, including genetic deletion or pharmacological block of OPC GABA_B_R decreased TWEAK expression (Fig. 3d, Extended Data Fig. 10).

To confirm that OPC-derived TWEAK can indeed induce neuronal apoptosis, we treated primary cortical neurons with conditioned medium obtained from OPC Oli-neu cultures (Fig. 3e). A selective TWEAKR antagonist, L524-0366 was added to the conditioned medium at 20 µM to block the binding of TWEAK on interneurons (Fig. 3e, f). While the conditioned medium from the OPC cell line increased interneuron apoptosis (Fig. 3g, h), co-treatment with the TWEAKR antagonist prevented the cell death. These results further substantiate that OPCs induces interneuron apoptosis via TWEAK secretion.

In summary, our results demonstrate that OPCs induce interneuron apoptosis by releasing TWEAK upon GABA_B_R activation. Blocking this pathway prevents the apoptosis of interneurons. The concomitant dysregulation of interneuron density results in a reduced inhibitory GABAergic tone and altered myelination in the mPFC.

### Interrupted OPC-interneuron communication after GABA_B_R loss in OPCs during development generates cognitive impairment in adulthood

The mPFC is responsible for cognitive processes and their regulatory finetuning is guaranteed by the E/I balance ^32^. To test whether the selectively reduced interneuron apoptosis and the associated structural myelin alterations can generate defects in neural circuits and cognition, we further performed *in vivo* investigations using electrophysiology and behavioral analysis in 9w old animals after gene ablation was induced at P7/8 (Fig. 4a).

**Figure 4.**
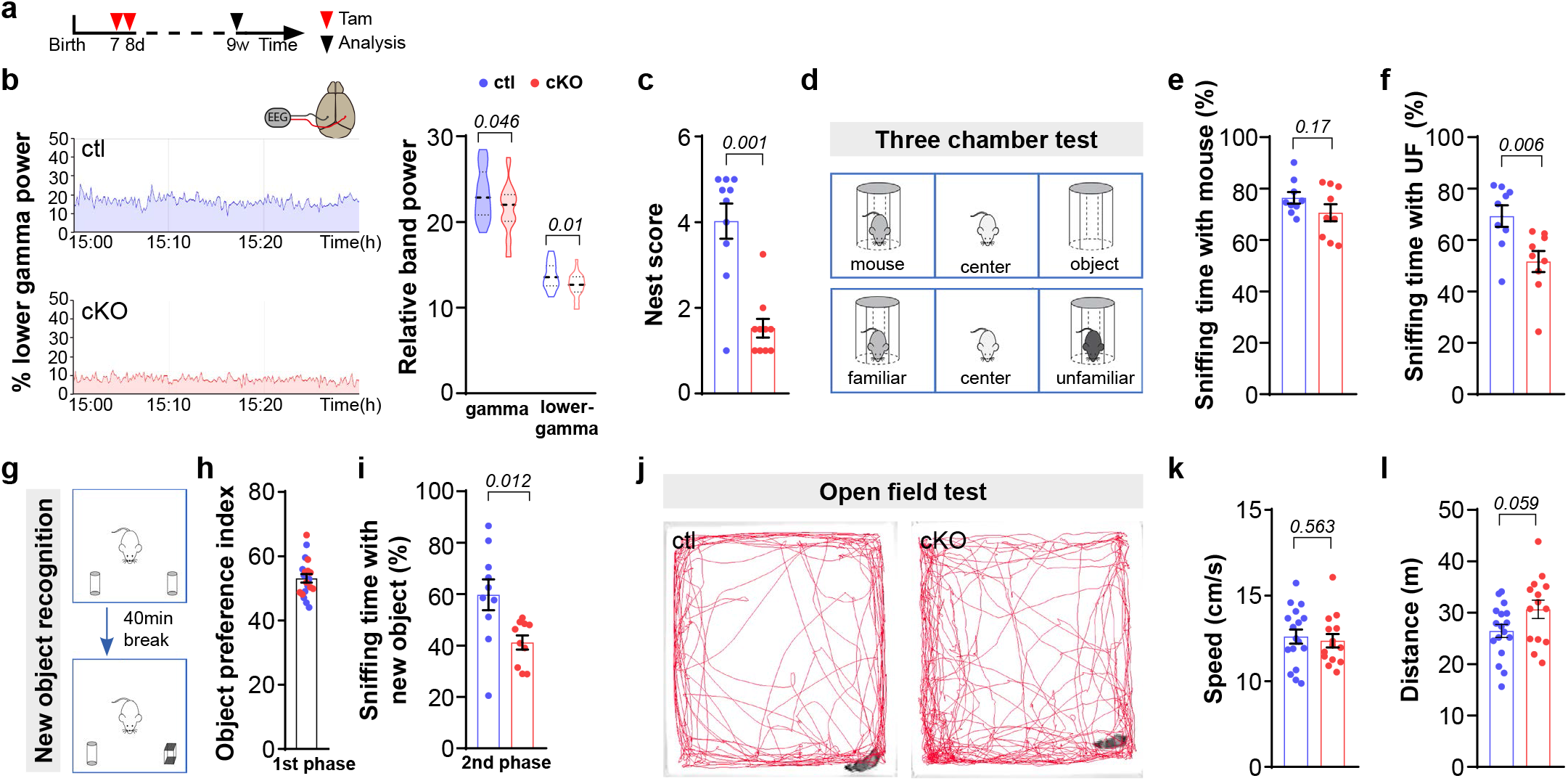
In OPC-GABA_B_R cKO mice perturbed electrical activity translates into deficits of social behavior. ***a***, Scheme of experimental plan. ***b***, Representative diagram of lower gamma band acquired by ECoG recordings. Violin plot shows decreased gamma oscillation in the cKO mouse brain. (*ctl=24 from 4 mice, cKO=24 from 4 mice, unpaired t-test*). ***c***, Impaired nest building ability of mutant mice. (*ctl=9, cKO=10, Mann Whitney test*). ***d***, Scheme of three-chamber behavior test. ***e***, Quantification of the sniffing time with mouse (***e***) or with unfamiliar mouse (***f***) among total sniffing time (*ctl=9, cKO=9, unpaired t-tests*). ***g***, Scheme of new object recognition test. ***h***, Assessment of object preference *(ctl=10, cKO=10)*. ***i***, Percentage of sniffing time with new object among total sniffing time. *(ctl=10, cKO=10, upaired t-test*). ***j***, Representative trajectory charts of the animals during the open field test. ***k, l***, Both ctl and cKO mice exhibited similar motor activities shown by the speed and distance analysis. (*ctl=10, cKO=14, unpaired t-tests)*.

The patch-clamp recordings described above had revealed a reduced firing rate of interneurons suggesting an imbalance of E/I in the mutant mPFC (Fig. 2c, d). Impaired cortical network activity of the mutant mice was also detected in electroencephalographical (EEG) recordings. The power of the gamma wave, especially the lower gamma wave band (30-50 Hz) was slightly, but significantly decreased by about 10% in cKO mice (Fig. 4b).

To evaluate the impact of circuit dysregulation in the mutant mPFC for the living animal, we challenged mice for their cognitive performance employing tasks of social cognitive behavior which had been associated with proper mPFC function, i.e. social novelty and new object recognition ^33^. In addition, nest building behavior is often introduced as a gauge of general well-being of the mice and also being an early sign of cognition decline ^34^. We observed that the mutant mice built significantly poorer nests with less than half score of the control animal nests, indicating putative cognition deficits of the mutant mice (Fig. 4c). The three-chamber test was used to assess social cognition by analyzing mice for a form of general sociability and their interest in social novelty (Fig. 4d). First, the three-chamber test was used to monitor general social behavior, i.e. preference of other mice over objects. Both, ctl and cKO mice exhibited more interest in the mouse rather than the object during the first test phase, indicating no disorder in sociability of the mutant mice (Fig. 4e, Extended Data Fig. 11). At the second phase, a social novelty preference was assessed. Control animals investigated the unfamiliar mouse more than the familiar one, while the cKO mice exhibited no novel social preference as they spent similar time in both chambers (Fig. 4f, Extended Data Fig. 11). Employing a new object recognition test (Fig. 4g), we again observed the cognition defect of cKO mice. Control animals were able to distinguish the new object replaced after a 40 min break from the habituation, while the cKO mice showed similar interest to both of the old and new objects (right panel in Fig. 4g-i). Open field test showed that the motor activity of the cKO mice (in terms of the speed and total distance) was not affected (Fig. 4j-l).

Hence, we conclude that GABA_B_Rs of OPCs are essential for cognition by modifying inhibitory circuits in the mPFC.

## Discussion

A balance of excitation and inhibition (E/I) in the medial prefrontal cortex (mPFC) is of key importance for mammalian cognition. Establishing correct cell densities and subsequent myelination have been identified as pivotal elements during CNS development ^6, 17^. This has been well documented for fast-spiking parvalbumin-positive (PV^+^) interneurons. Here, we highlight OPCs as indispensable regulators of interneuron population and myelination in the mPFC by providing a molecular explanation and demonstrating how impaired bidirectional OPC-interneuron signaling affects social cognitive behavior. OPCs at the first two postnatal weeks release TWEAK to regulate interneuron apoptosis. Genetic ablation of GABA_B_R in OPCs at the first postnatal week promoted interneuron survival. However, these neurons displayed hypoactivity, less contacts with OPCs during development and deteriorated myelin structures in the adulthood. Notably, these mutant mice exhibited severe cognitive defects in their social behavior.

During development, large populations of immature interneurons invade the cortex where their cell density is adjusted by programmed cell death ^9, 10^. For assuring their survival, these neurons form connections with adjacent cells and receive retrograde signals from their targets ^10, 11^. Interneuron apoptosis drastically increases at p7 ^7^, shortly after formation of interneuron-OPC synapses at p4-5 ^12^. Based on this established knowledge, we hypothesized that OPCs could exert an apoptotic impact to their presynaptic partners. Indeed, our data provide strong evidence that, starting from postnatal day 5 on, OPCs elicit an apoptotic cascade of adjacent interneurons to adjust their density. In OPCs, the regulatory process employs a GABA_B_R-TWEAK signaling pathway. TWEAK, TNF super family member 12 (TNFSF12), is TNF-like weak apoptotic factor. Membrane-bound and cleaved isoforms of TWEAK are both able to induce cell apoptosis by binding to the TWEAK receptor (TWEAKR, also known as FN14) on the neuronal side ^31, 35, 36^. According to our data GABA_B_R-TWEAK signaling preferentially affects interneurons, suggesting this ‘kill me’ signal acting highly localized. Indeed, TWEAK is specifically recruited to synapses where TWEAKRs are expressed ^37^. OPCs directly sense GABAergic signals at their interneuron-OPC synapse. Subsequent signaling triggers TWEAK expression and translocation to the OPC surface, where it could directly bind interneuronal TWEAKRs. Alternatively, the TWEAK ectodomain could be proteolytically cleaved and act as a soluble factor. However, this scenario could also happen at soma-somatic contact sites between OPCs and interneurons ^38, 39^. The regulatory impact appears to be highly specific since the OPC-induced interneuron apoptosis was restricted to the first two postnatal weeks, a well-established time window of interneuron apoptosis ^7, 30^. This also explains why the cKO induction at the age of 4w did not impact OPC differentiation and myelination, although GABA_B_R expression peaks in OPCs at 4w (data not shown). Please note, instead of a gradual decline, we found a slight increase of PV^+^CC3^+^ cells at p14 compared to p10, probably due to the age-dependent increase of PV expression described for fast spiking interneurons ^40^.

Neuronal overpopulation is often accompanied with suppressed activity ^6, 8^. In the OPC-GABA_B_R cKO mice, we observed excessive number of PV^+^ interneurons (about 30 % increase) while attenuated vGAT puncta (reduced to 80 %), suggesting single PV^+^ interneuron exhibits suppressed activity (about 60% ≈ 80 %÷130 %). This was also shown by the sIPSP recording which was decreased to 60 % in the cKO mPFC. Differentiation of oligodendrocytes and subsequent myelination are strongly affected by neuronal activity ^22, 26, 41^. Indeed, we observed a suppression of the inhibitory tone in the mPFC at p10 followed by a change in oligodendrocyte density detected at p14. Concomitantly, the oligodendrocytes formed less myelin (only 40 % of control scenario) around the interneuronal axons and altered paranodal structures in the adult mPFC which are essential for precise action potential propagation. Such reduction is attributed to the restrained PV^+^ neuron activity (60 %) and oligodendrocyte population (74 %) (60 %×74 % ≈ 40 %). Regardless, the morphological deterioration was only observed in the mPFC, but not in the corpus callosum composed axons of excitatory neurons only ^21^. In line with these neuron-type specific observations, the qRT-PCR results indicated reduced levels of MBP, but not PLP mRNA expression in the mutant mPFC, both at p14 and 9w of age. Axons of interneurons and excitatory neurons are different in respect to the protein content of their myelin sheaths. MBP is preferentially expressed in the myelin of inhibitory axons, while PLP is more prominent in excitatory axons PLP ^16^.

An early loss of the GABA_A_R γ2 subunit in OPCs reduced the firing rate of presynaptic PV^+^ interneurons and caused a similar myelin defect of longer nodal structures in PV^+^ neurons ^4^, the initially very different signal pathways of both, GABA_A_R and GABA_B_R, have to merge to affect myelin gene expression and OPC-axon contacts similarly. The GABA_A_R signaling between interneurons and OPCs is relatively well established ^4, 15, 42, 43^. E.g., the γ2 subunit of GABA_A_R contributes to the maintenance of the OPC, but has no impact on oligodendrocyte formation ^15^. However, the role of GABA_B_Rs for OPC differentiation is complex. We observed that GABA_B_R was essential for oligodendrogenesis at the early development but not at later stages. Similary, an *in vitro* study showed that activation of GABA_B_R by the selective agonist baclofen could stimulate the differentiation of cortical OPCs prepared from p0-2 pups ^14^. Since OPCs are heterogeneous with age ^44^, GABA_B_R of OPCs could contribute to the developmental diversity.

Obviously, the GABAergic signaling of OPCs plays pivotal roles for the myelination of interneurons, which ensures precise finetuning of the local neural circuitry ^45^. As seen in chandelier cells ^6^, excessive number of interneurons in the OPC-GABA_B_R cKO mice exhibited suppressed activity, and reduced myelination in the mPFC. All these physiological and morphological abnormalities certainly contributed to cognition impairment. A proper E/I ratio in the prefrontal cortex, especially early E/I balance in mPFC, is extremely important for social cognition ^32, 46^. When the pyramidal neuron activity was enhanced in the mPFC at early p7-11 caused a severe cognitive dysfunction in the adulthood ^46^. In addition, despite relatively short axons, myelination of cortical interneurons (mainly PV^+^ neurons ^16^) contributes significantly to the finetuning of local activity in the mPFC and cortex, insuring proper behavior performance ^4, 6, 17^.

In conclusion, our study demonstrates that OPCs regulate inhibition in the mPFC via GABA_B_R/TWEAK signaling. During development, OPCs determine the density of interneurons by adjusting their apoptosis. Subsequently, correct myelination of interneurons, as an essential component of the network activity in the mPFC, determines the cognition and social behavior.

## Acknowledgement

We thank Daniel Schauenburg for excellent animal husbandry and Frank Rhode, Alexander Grissmer and Samantha Bechet for experimental assistance. We are also grateful to Prof. Jaqueline Trotter (University of Mainz, Mainz, Germany) for providing the Oli-neu cell line and Prof. Leda Dimou (University of Ulm, Ulm, Germany) for sharing MACs sorting protocols of OPCs. This work was supported by grants from the Deutsche Forschungsgemeinschaft (SPP 1757, SFB 894 to FK; FOR2289 to AS, FK, SPP1757 Young Investigator grant to XB), EraNet-Neuron BrIE to FK and HOMFORexzellent2017 program to XB.

## Author contributions

XB initiated the project. LF, XB, NZ, LC, NH, CL and AS performed experiments. LF, XB, NZ, LC and RZ analyzed data. BB, WH. HC and CM provided materials. XB and FK designed and supervised the study and wrote the manuscript with comments of the other authors.

## Supplementary figure legends

**Supplementary Fig. 1.**
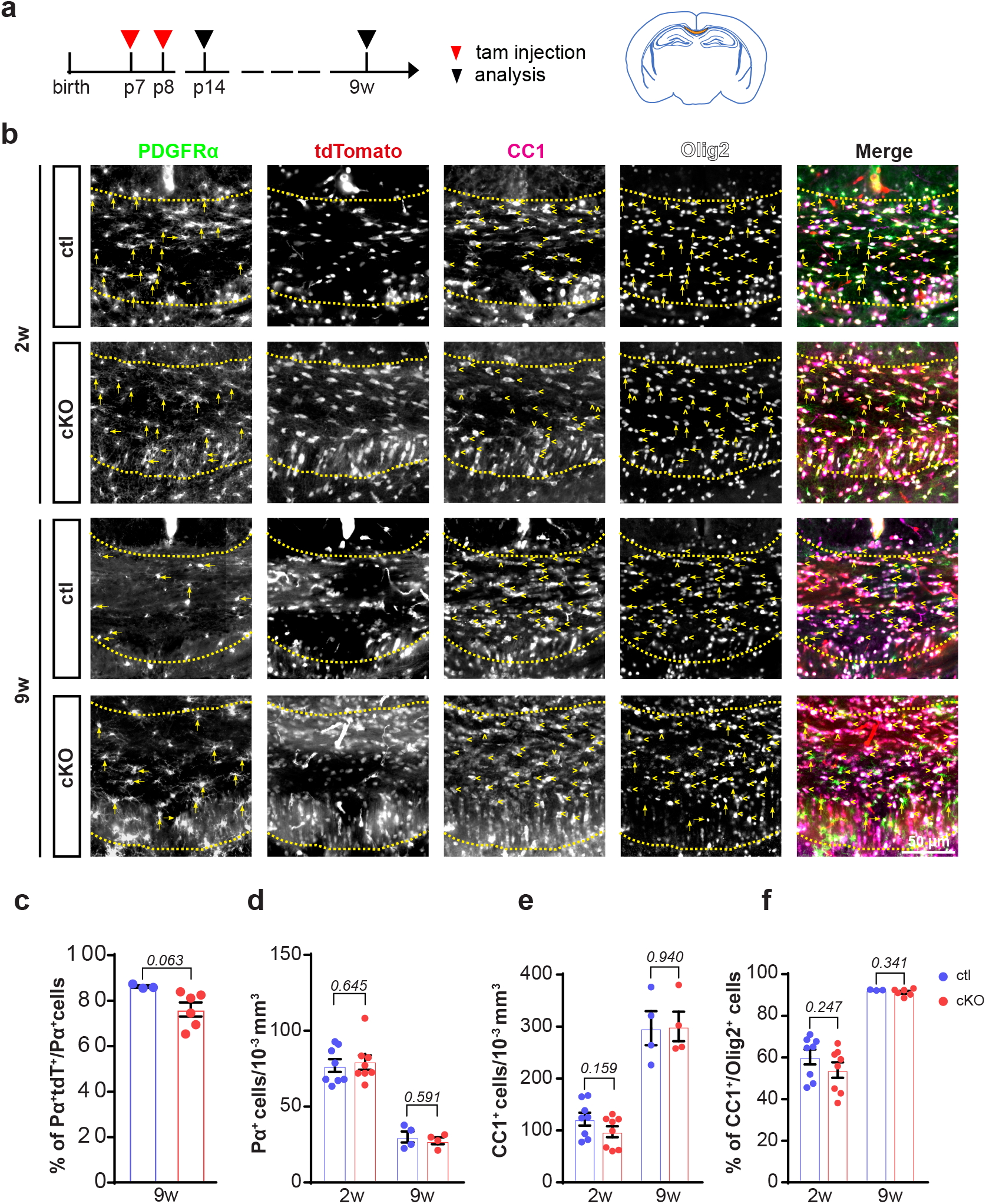
OPC and oligodendrocyte density is not altered in the corpus callosum of mutant mice. **a,** Scheme of experimental schedule (left panel) and analyzed brain region in highlight, i.e. corpus callosum (cc, right panel). **b,** Corpus callosal OPCs and oligodendrocytes of ctl and cKO mice were immunolabeled with PDGFRα and CC1 respectively at the age of 2w and 9w. Olig2 was used as lineage marker and tdTomato was indicator of recombined cells. **c,** OPC recombination efficiency in ctl and cKO cc (*ctl=86.22±0.57 (n=3), cKO=76.17±3.10 (n=6*), unpaired t-test). **d, e,** Cell density of OPCs (PDGFRα^+^) and oligodendrocytes (CC1^+^) in cc of 2w- and 9w-old mice were not changed. OPC: (*2w-ctl=77.02±4.21 (n=8), 2w-cKO=80±4.72 (n=8), unpaired t-test*) (*9w-ctl=30.01±3.68 (n=4), 9w-cKO=27.56±2.26 (n=4), unpaired t-test*); oligodendrocytes (*2w-ctl=121.70±12.31 (n=8), 2w-cKO=97.62±10.54 (n=8), unpaired t-test*) (*9w-ctl=296.90±32.83 (n=4), 9w-cKO=300.30±28.47 (n=4), unpaired t-test*). **f,** Percentage of oligodendrocytes among total Olig2^+^ cell population, considered as OPC differentiation rate, was not changed at both ages (*2w-ctl=60.26±3.61(n=8), 2w-cKO=53.98±3.73 (n=8), unpaired t-test*) (*9w-ctl=92.37±0.12 (n=3), 9w-cKO=91.33±0.69 (n=6), unpaired t-test*).

**Supplementary Fig. 2.**
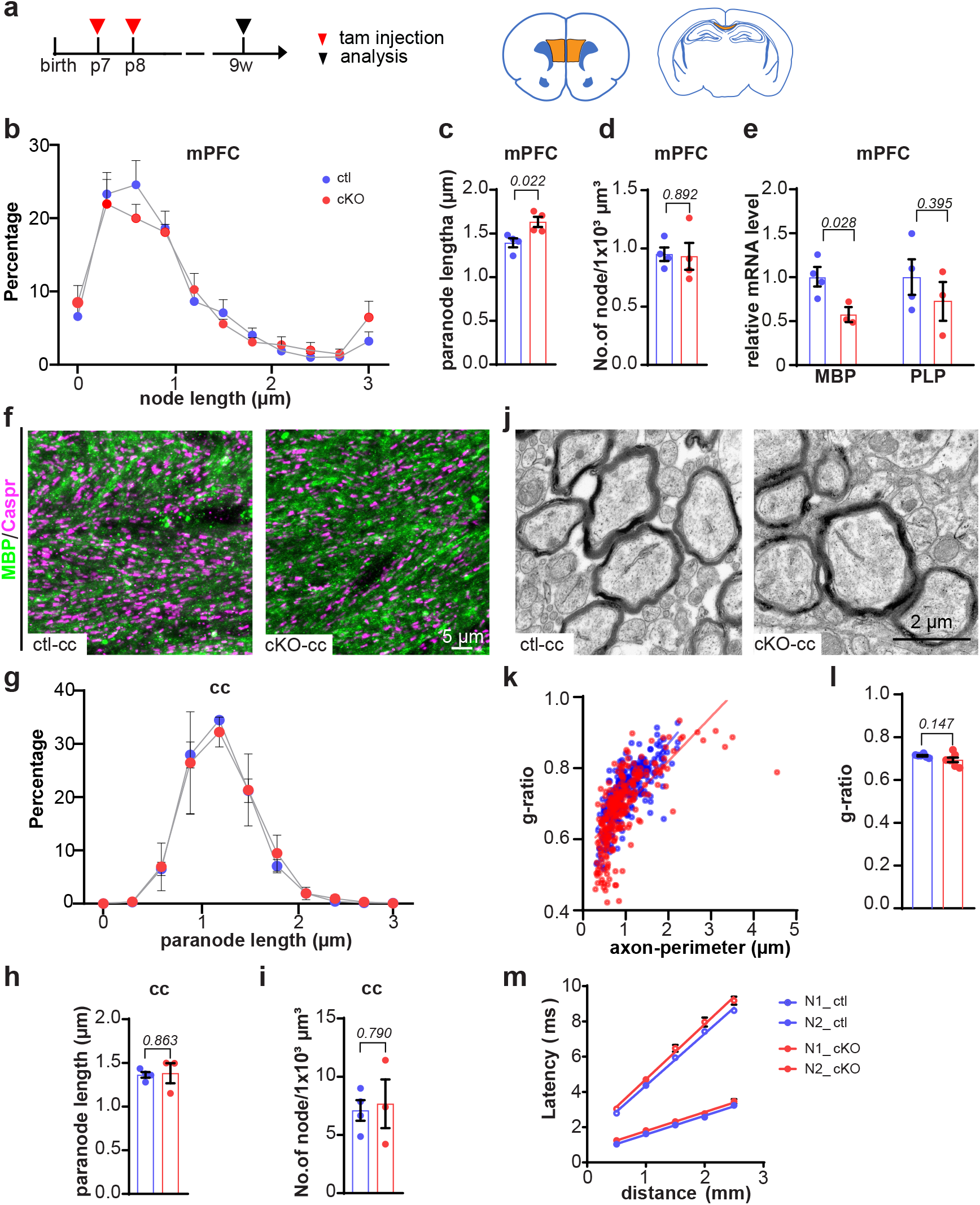
Parameters of myelination remain largely unaltered in the corpus callosum of mutant mice. **a,** Scheme of experimental schedule (left panel) and analyzed brain region in highlight, i.e. mPFC and cc. **b,** Quantification of node length of ctl and cKO mPFC. **c,** Paranode length is increased in the cKO mPFC. **d,** Density of nodes remained same in the cKO mPFC. **e,** Relative mRNA level of MBP is reduced in the cKO mPFC, while PLP mRNA is not affected (*ctl: n=4, cKO: n=3, unpaired t-test*). **f,** Immunostaining of Caspr and MBP in cc of ctl and cKO mice. **g-i,** Paranodal length and node density in ctl and cKO cc remained same. **j,** Electron microscopic analysis of myelin ultrastructure in sagittal section of cc from ctl and cKO mice. **k, l,** Quantification of g-ratios of axonal myelination shows unchanged myelination in cKO cc. **m,** Compound action potential conduction velocity was not altered indicated by patch clamp recordings. N1 indicates non-myelinated axonal conduction velocity, while N2 is for myelinated axons.

**Supplementary Fig. 3.**
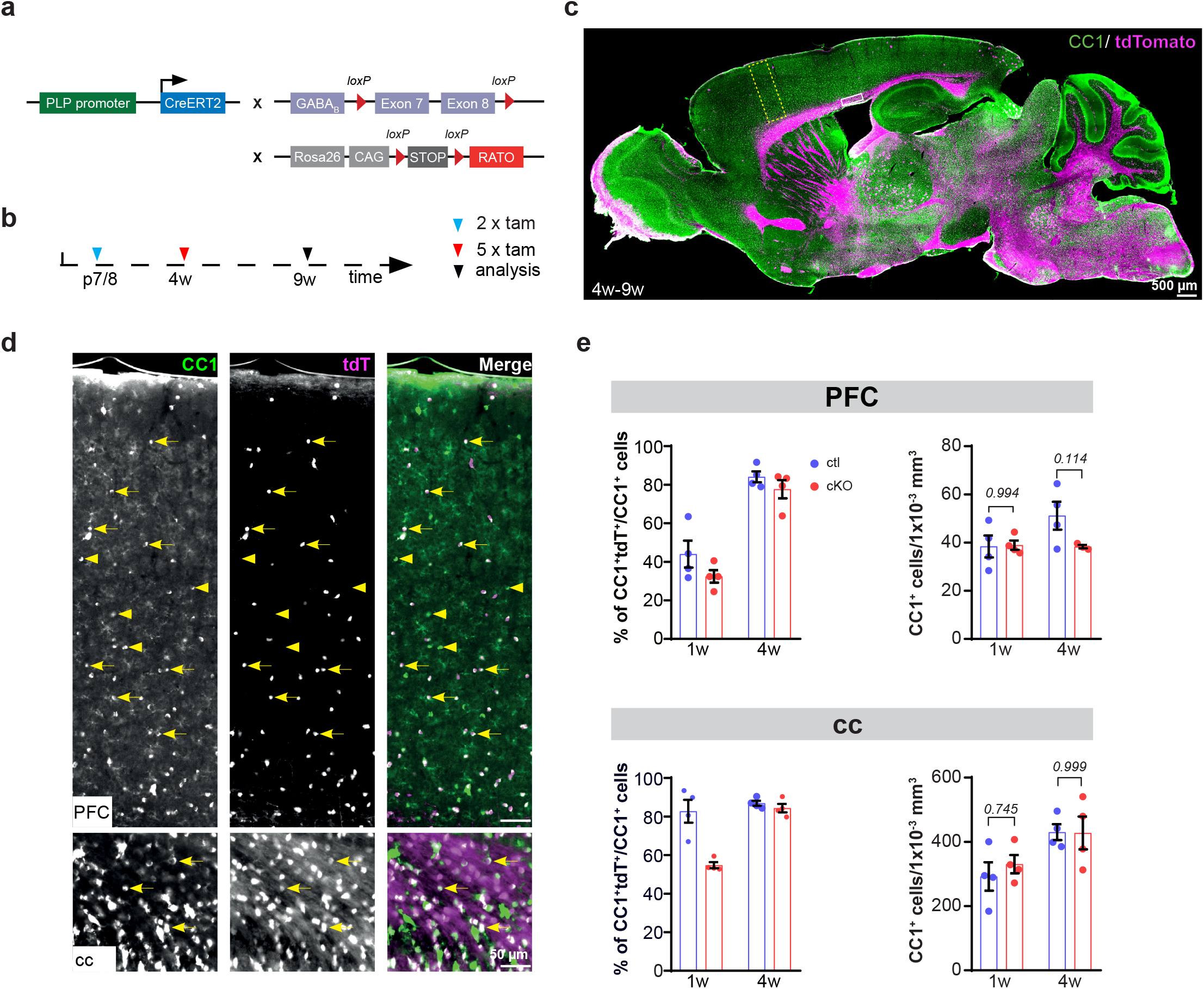
Ablation of GABA_B_R in oligodendrocytes do not affect formation and density of oligodendrocytes. **a,** Scheme of transgene construction. **b,** Experimental schedule. **c,** Overview of CC1 immunostaining in a sagittal section of 9w old ctl mouse. Extensive tdTomato distribution shows a high recombination in different brain regions of transgenic mouse. **d,** Magnified images from prefrontal cortex (yellow box in **c**) and corpus callosum (white box in **c**). **e,** Quantification of oligodendrocyte recombination efficiency in PFC were identical, while cKO mice showed smaller recombination efficiency in cc. However, density of oligodendrocyte cells did not differ between ctl and cKO mice in both PFC and cc. arrows: recombined oligodendrocyte; triangles: non-recombined oligodendrocyte.

**Supplementary Fig. 4.**
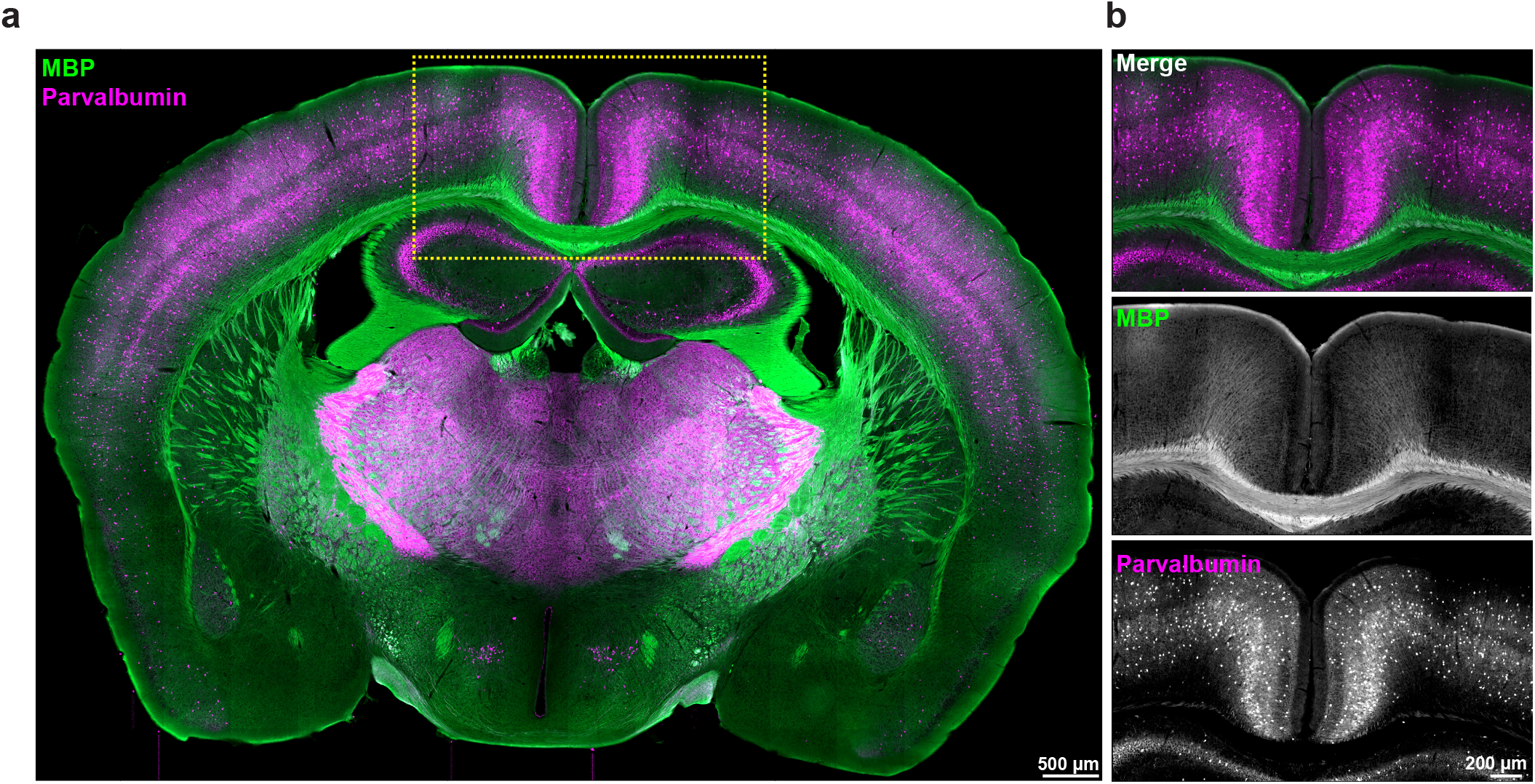
Dense population of parvalbumin-expressing interneurons in the adult mouse cortex. **a,** Overview of coronal brain slices immunostained with parvalbumin and MBP shows abundant PV^+^ neurons in the somatosensory cortex. **b,** Magnified view (yellow box in **a**) shows the absence of PV^+^ axons in the corpus callosum.

**Supplementary Fig. 5.**
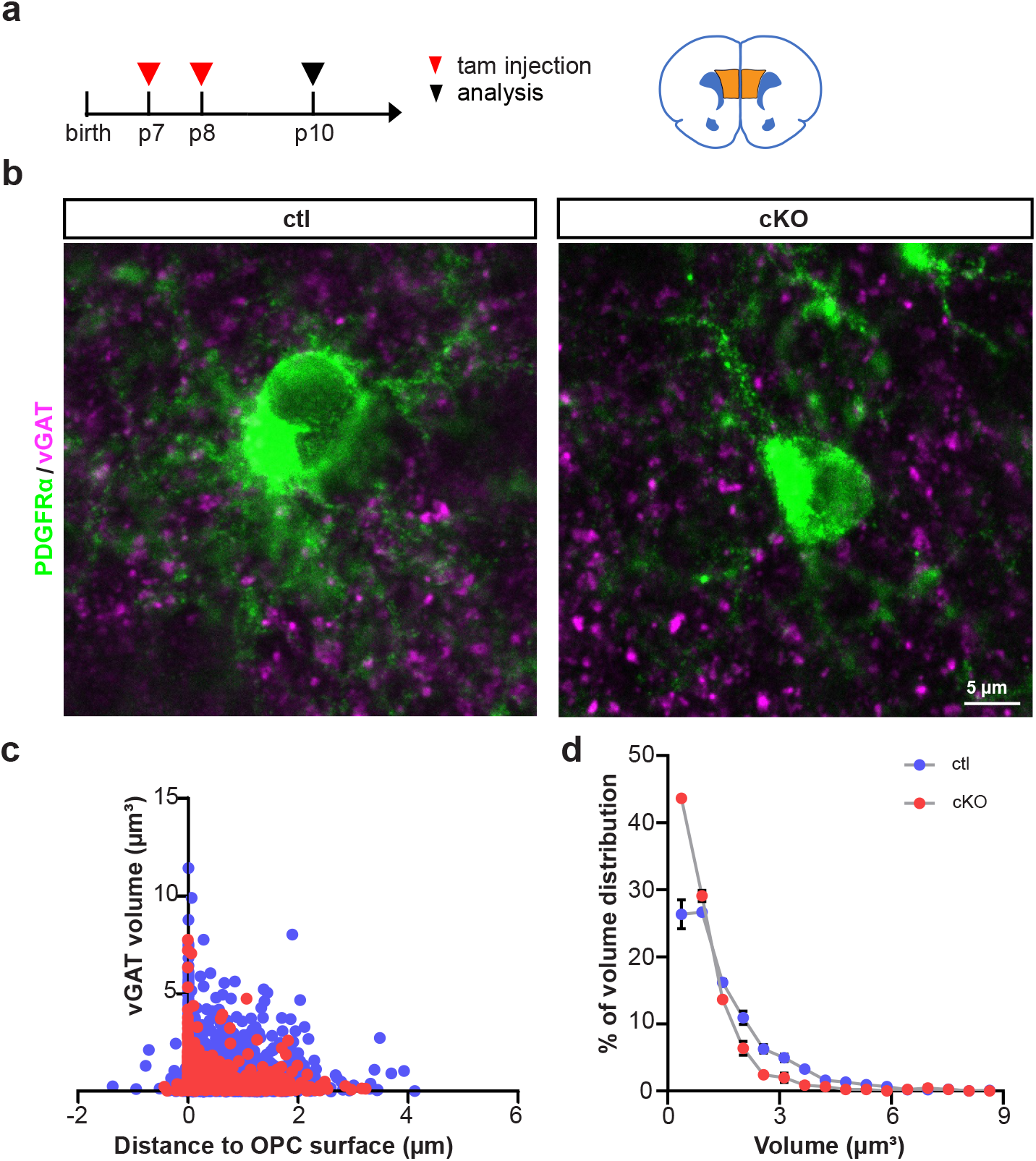
At the onset of myelination, GABA_B_R-deficient OPCs receive less GABAergic input in the mPFC mutant mice. **a,** Experimental timeline and analyzed brain region (mPFC) in highlight.**b,** Immunostaining of PDGFRα and vGAT in the mPFC of p10 mice. **c,** Quantitative analysis of vGAT volume and the distance to OPC surface (*ctl: n=4, cKO: n=4*). **d,** The volume of vGAT in cKO mPFC was smaller than that in ctl group.

**Supplementary Fig. 6.**
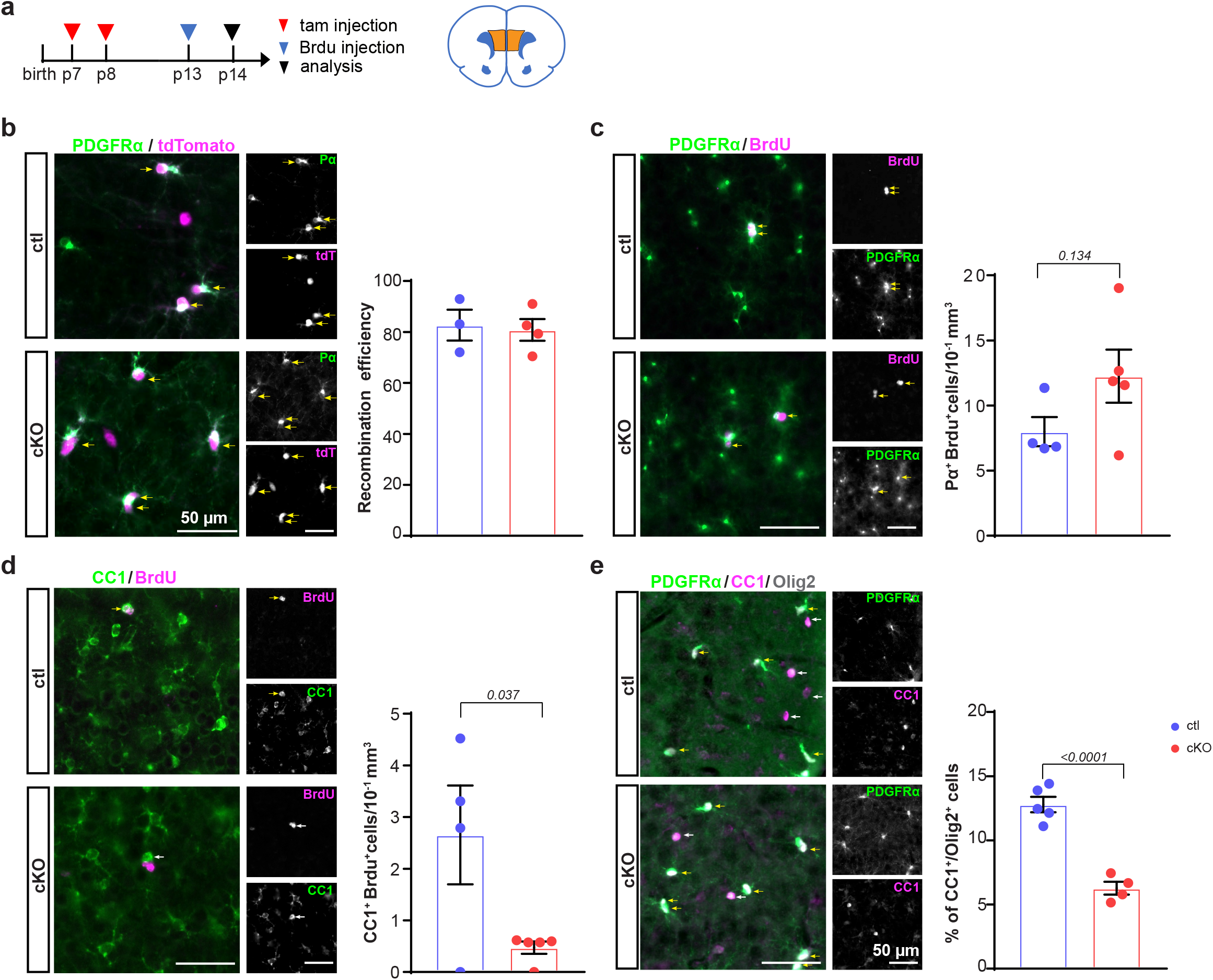
The differentiation of GABA_B_R-deficient OPCs is suppressed in the mutant mPFC. **a,** Scheme of experimental schedule and analyzed brain region (mPFC). **b,** Recombination efficiency of OPCs (% of Pα^+^tdt^+^/Pα^+^) at p14 is identical in the ctl and cKO mPFC. **c,** Immunostaining and quantification of OPCs incorporated with BrdU shows no change in OPC proliferation in the cKO mPFC at p14 (*ctl: n=4, cKO: n=5, unpaired t-test*). **d,** Immunostaining and quantification of oligodendrocytes incorporated with BrdU shows decreased OPC differentiation in the cKO mPFC at p14 (*ctl: n=4, cKO: n=5, unpaired t-test*). **e,** The portion of oligodendrocytes among the oligodendrocyte lineage declined in cKO mPFC at p14 (*ctl: n=5, cKO: n=4, unpaired t-test*).

**Supplementary Fig. 7.**
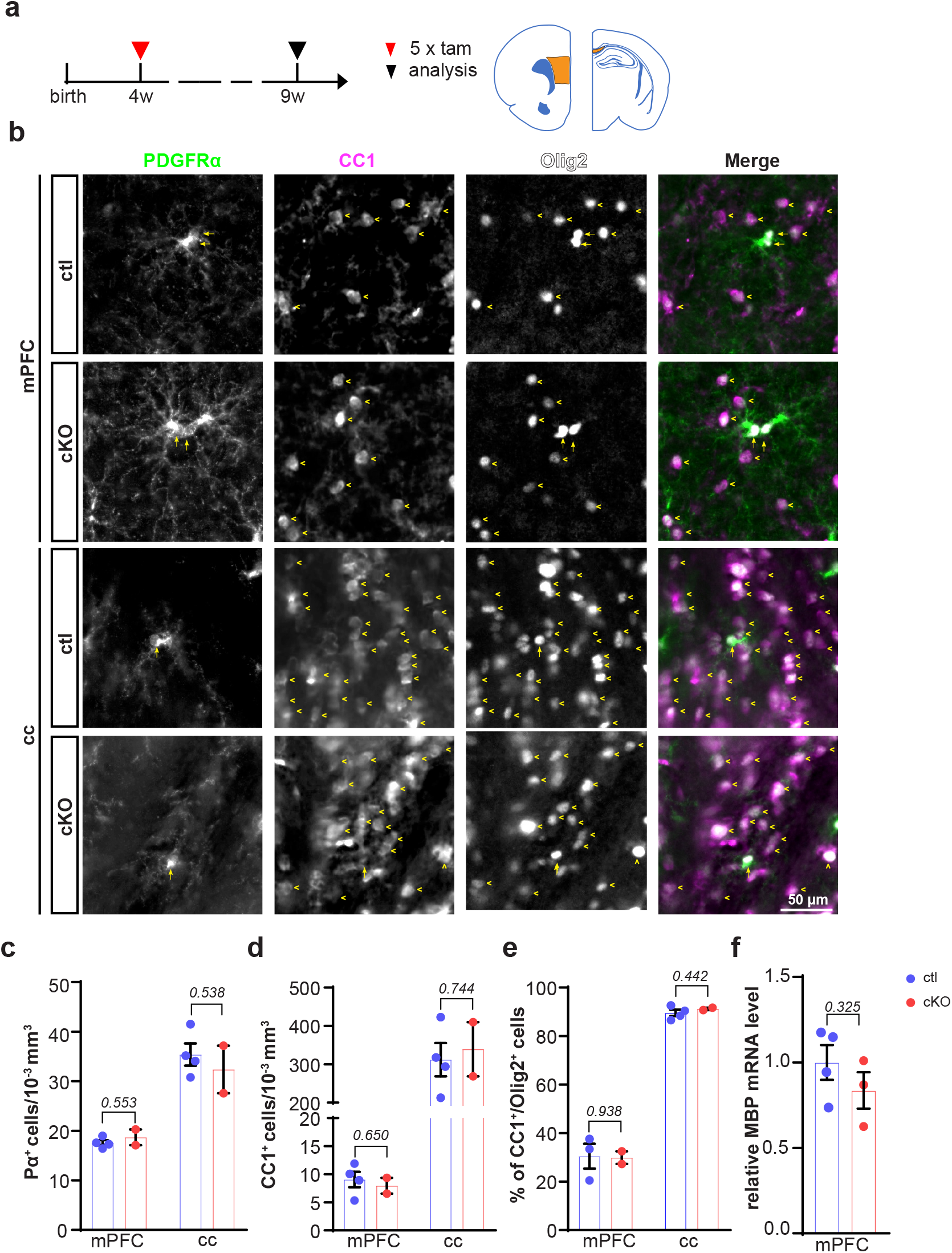
Controlled ablation of GABA_B_R in OPCs of young adult mice does not affect oligodendrogenesis. **a,** Scheme of experimental schedule and the analyzed brain regions (mPFC and cc). **b,** Immunostaining of OPCs (PDGFRα) and oligodendrocyte (CC1) with Olig2 (lineage marker) at ctl and cKO mouse mPFC and cc at the age of 9w. **c, d,** Cell densities of OPCs (PDGFRα^+^) and oligodendrocytes (CC1^+^) of cKO animals were the same as those for ctl mice in both mPFC and cc. mPFC. *(PDGFRα^+^: (ctl=17.58±0.53 (n=4), cKO=18.69±1.58 (n=2), unpaired t-test)); (CC1^+^: (ctl=9.05±1.38 (n=4), cKO=7.95±1.40 (n=2), unpaired t-test));* cc: *(PDGFRα^+^: (ctl=35.38±2.25 (n=4), cKO=32.37±4.81 (n=2), unpaired t-test)); (CC1^+^: (ctl=311.81±43.39 (n=4), cKO=339.20±70.53 (n=2), unpaired t-test))*. **e,** The proportion of oligodendrocyte among total lineage cells did not change, suggesting OPC differentiation remained same in the mutant mouse mPFC and cc. mPFC: *(ctl=30.49±5.13 (n=3), cKO=29.91±2.62 (n=2), unpaired t-test); cc: (ctl=89.52±1.30 (n=3), cKO=91.21±0.471 (n=2), unpaired t-test)*. **f,** MBP expression is not altered in the cKO mPFC at mRNA level (*ctl: n=4, cKO: n=3, unpaired t-test*).

**Supplementary Fig. 8.**
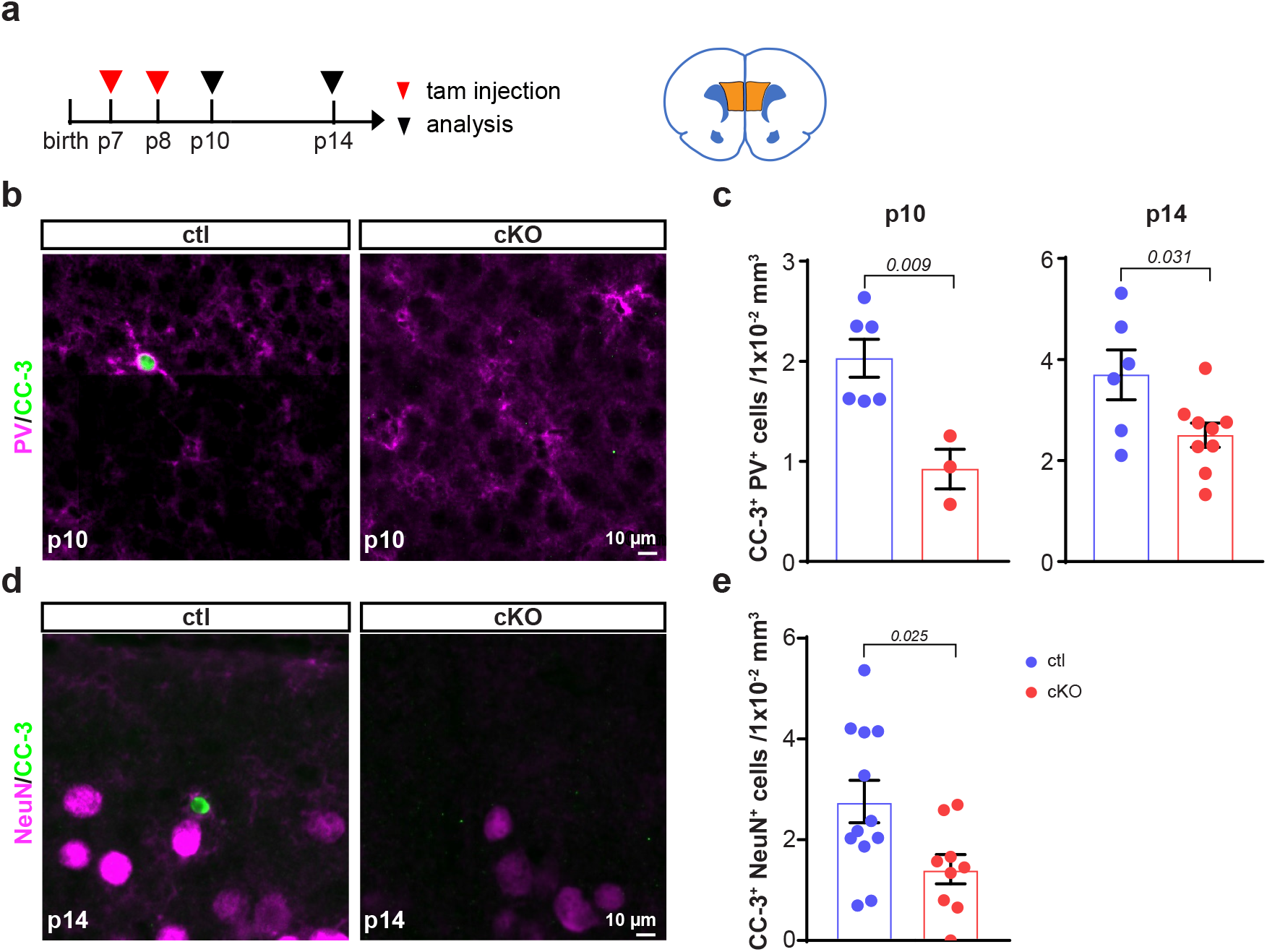
The apoptosis of interneurons is mitigated in the mPFC of mutant mice at p14. **a,** Timeline of experiments and scheme of analyzed brain region mPFC in highlight. **b,** Immunostaining of PV^+^ neurons with apoptotic marker (cleaved caspase-3) at p10 of ctl and cKO mPFC. **c,** Quantification of apoptotic PV^+^ neurons at p10 and p14 shows reduced PV^+^ neuronal apoptosis in the cKO mPFC compared to ctl (p10*: ctl=2.03±0.47 (n=6), cKO=0.92±0.34 (n=3), unpaired t-test;* p14: *ctl=3.70±1.21 (n=6), cKO=2.50±0.72 (n=9), unpaired t-test)*. **d,** Representative images of apoptotic neurons at p14 of ctl and cKO mPFC. **e,** Quantification of apoptotic total neurons (NeuN^+^CC-3^+^) at p14 *(ctl=2.76±0.42 (n=12), cKO=1.42±0.29 (n=9), unpaired t-test)*.

**Supplementary Fig. 9.**
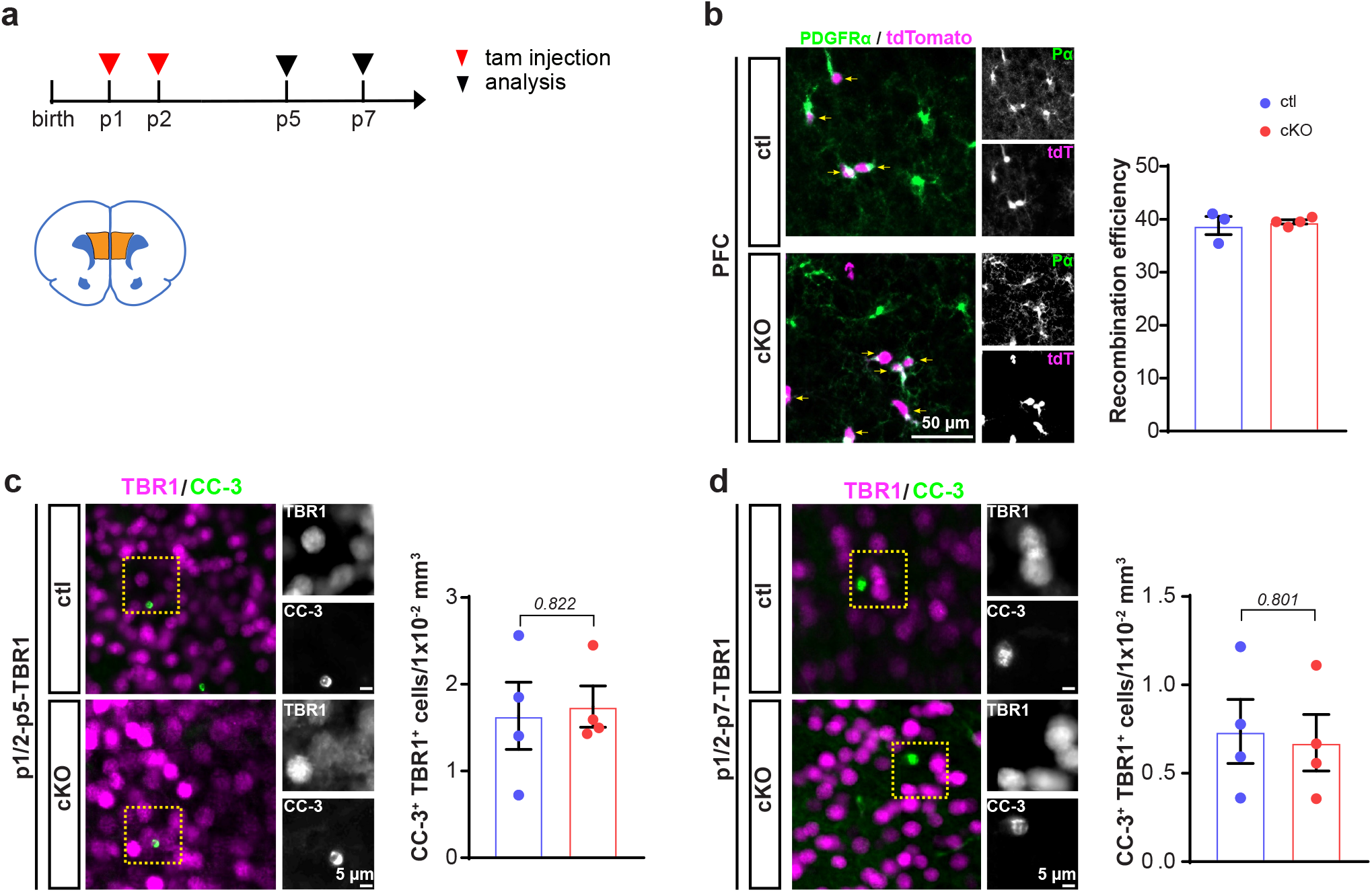
Ablation of GABA_B_R expression in OPCs does not affect the apoptosis or survival of excitatory neurons. **a,** Scheme of experimental schedule (left panel) and analyzed brain region in highlight. **b,** About 40 % of OPCs are recombined (% of PDGFRα^+^tdT^+^/PDGFRα^+^) at p7 both in the ctl and cKO mPFC. **c, d,** Immunostainings and quantification of excitatory neurons (TBR1^+^) with apoptotic marker CC-3 in mPFC of ctl and cKO pups at p5 (**c**) and p7 (**d**) (p5: *ctl=1.64±0.39 (n=4), cKO=1.74±0.24 (n=4), unpaired t-test;* p10*: ctl=0.74±0.18 (n=4), cKO=0.67±0.16 (n=4), unpaired t-test*).

**Supplementary Fig. 10.**
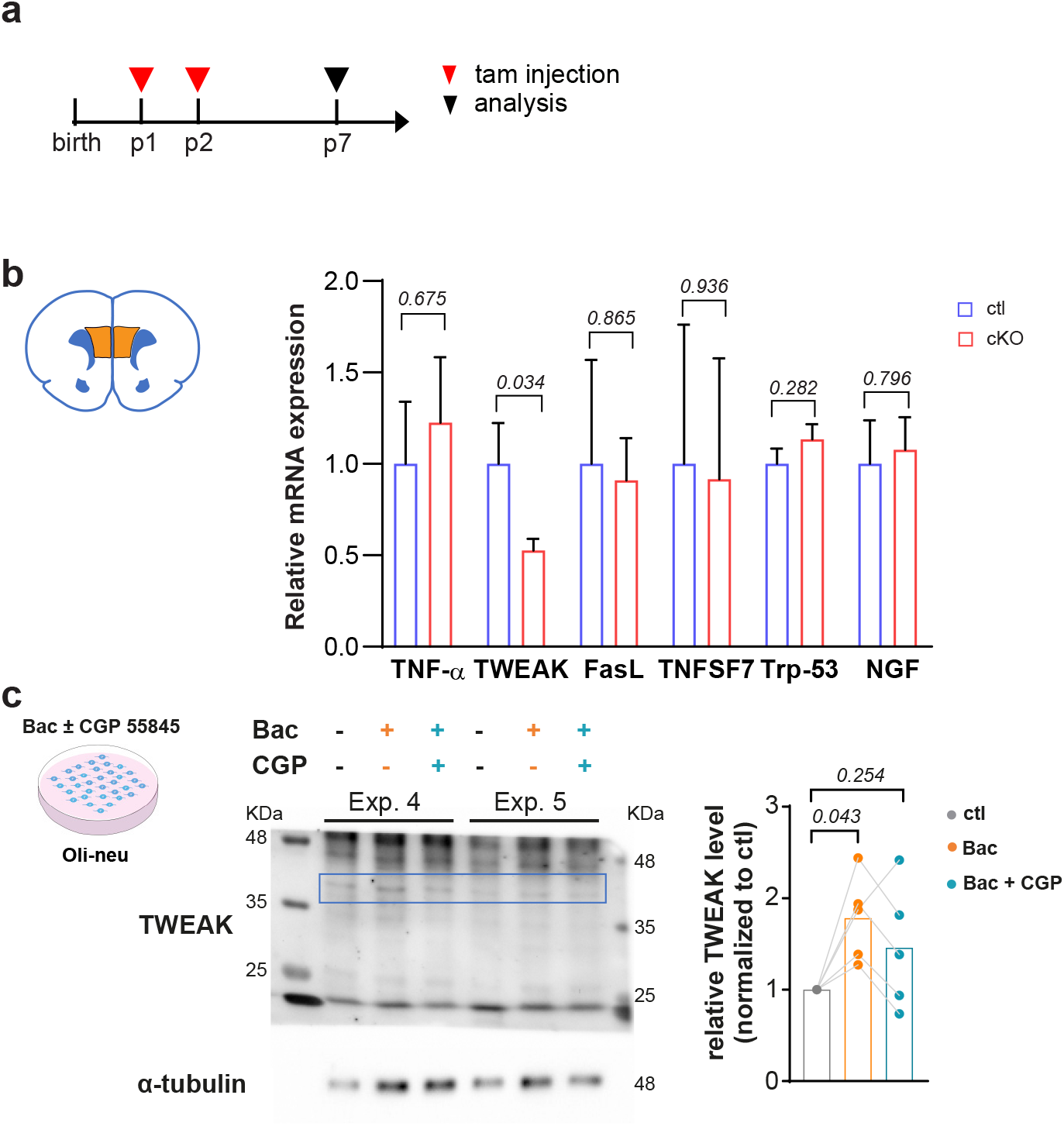
TWEAK expression in OPCs is regulated by GABA_B_Rs. **a,** Scheme of experimental design. **b,** Relative mRNA level of cytokines in the mPFC of ctl and cKO mouse at p7. *(TNF-α: ctl=1±0.34 (n=4), cKO=1.23±0.36 (n=6), unpaired t-test); (TWEAK: ctl=1±0.10 (n=4), cKO=0.67±0.08 (n=6), unpaired t-test); (FasL: ctl=1±0.57 (n=2), cKO=0.91±0.23 (n=4), unpaired t-test); (TNFSF7: ctl=1±1.00 (n=4), cKO=0.93±0.67 (n=6), unpaired t-test); (Trp53: ctl=1±0.11 (n=4), cKO=1.14±0.08 (n=6), unpaired t-test); (NGF: ctl=1±0.24 (n=5), cKO=1.08±0.18 (n=6), unpaired t-test)*. **c,** Western blot analysis of TWEAK expression in Oli-neu cells treated with 50 µM Baclofen at the presence or absence of 20 µM CGP 55845 (n=5 independent experiments). The isoform in 36 KDa (blue box) was analyzed.

**Supplementary Fig. 11.**
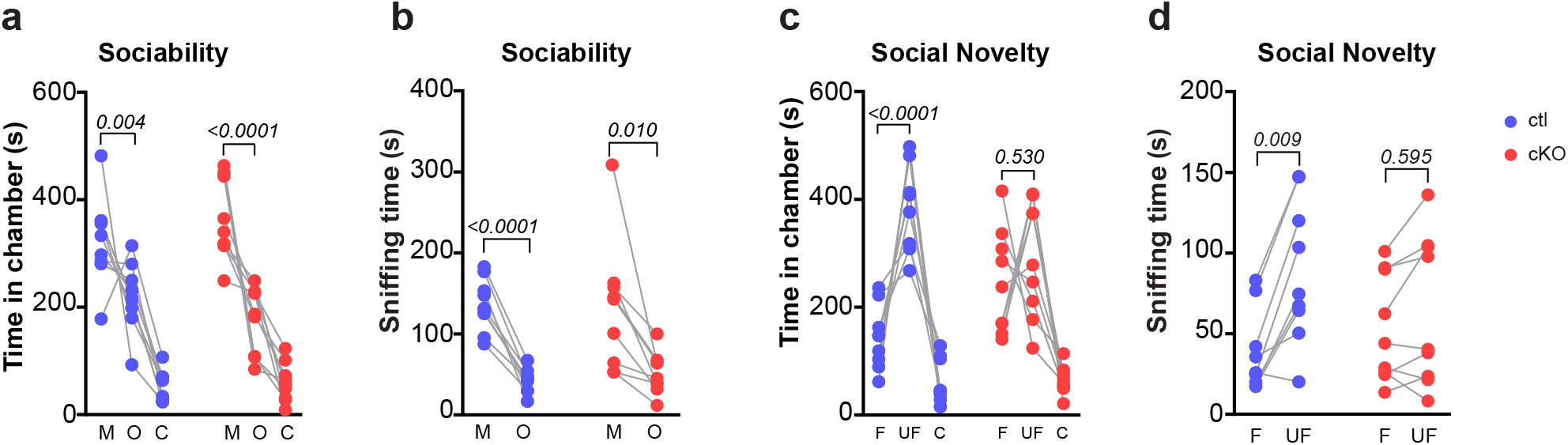
OPC-GABA_B_R cKO mice exhibit impaired social behavior. **a, b,** Mutant mice stayed longer at the chamber of mouse than object, indicating that sociability of the mutant mouse is not affected. **c, d,** Mutant mice spent similar time at the chambers of familiar and unfamiliar mouse, suggesting impaired cognition of the cKO mice (*ctl (n=9), cKO (n=9), paired t-test*).

## Supplementary information for

### Materials and Methods

#### Ethics statement

Animal husbandry and procedures were performed at the animal facility of CIPMM, University of Saarland according to European and German guidelines for the welfare of experimental animals. Animal experiments were approved by the Saarland state’s ‘‘Landesamt für Gesundheit und Verbraucherschutz” in Saarbrücken/Germany (animal license number: 65/2013, 12/2014, 34/2016, 36/2016 and 08/2021).

#### Animals

To conditionally knock out (cKO) GABA_B1_ receptors in oligodendrocyte precursor cells (OPCs), TgH(NG2-Cre^ERT2^)^1^ mice were crossbred to GABA_B1_^lox511/lox511^ mice (flanking exon 7 and 8 of *gabbr1*)^2^. Mice with genotypes of NG2^ct2/wt^ x GABA_B1_R^fl/fl^ were used as cKO and the littermates NG2^wt/wt^ x GABA_B1_R^fl/fl^ or NG2^ct2/wt^ x GABA_B1_R^wt/wt^ were controls after tamoxifen application. To visualize the recombined cells, we crossbred the double transgenic mice with TgH(Rosa26-CAG-^fl^STOP^fl^-tdTomato) (Rosa26-tdTomato) ^3^.

For oligodendrocyte specific deletion of GABA_B1_R, we took advantage of TgN mice (PLP-Cre^ERT2^)^4^ crossbred with GABA_B1_^lox511/lox511^. Similarly, PLP^ct2/wt^ x GABA_B1_R^fl/fl^ mice were used as cKO, while PLP^wt/wt^ x GABA_B1_R^fl/fl^ or PLP^ct2/wt^ x GABA_B1_R^wt/wt^ were controls after tamoxifen application. Mouse ages are indicated in the main text and figure legends. Behavioral tests were carried out only with male mice at the age of 9 weeks, while both genders were used in the other experiments.

#### Tamoxifen administration

Tamoxifen was dissolved in Miglyol (3274, Caesar & Loretz GmbH) to a final concentration of 10 mg/ml. Tamoxifen was intraperitoneally injected to the mice depending on the body weight (100 mg/kg body weight). The time points of injections are indicated in the figures. Only for the pups treated at postnatal day 1 (p1) and 2, tamoxifen was injected to the lactating mother with the same protocol ^5^. For 4-week-old mice, tamoxifen was injected once per day for five consecutive days ^6^.

#### Immunohistochemistry

Mice were perfused with PBS and 4 % paraformaldehyde (PFA). Mouse brains were post fixed with 4 % PFA at 4°C overnight. Free floating brain slices (40 µm thickness) were cut in coronal or sagittal sections using a Leica VT1000S vibratome. For immunocytochemistry, cells on coverslips were fixed with ice cold 4 % PFA for 15 min. Slices or coverslips were incubated with blocking solution containing 5 % horse serum and 0.5 % Triton X-100 at room temperature (RT) for 1 hour (hr), followed by primary antibody incubation at 4 °C overnight and secondary antibody incubation at RT for 2 hrs in blocking solution. Primary and secondary antibodies are listed in the **Supplementary table 1 and 2,** respectively.

#### Magnetic cell separation (MACS) of OPCs

MACS sorting of OPCs was performed according to the manufacturer’s instruction (Miltenyi Biotec) with some modifications. Mice were perfused with cold Hank’s balanced salt solution without calcium and magnesium (HBSS, H6648, Gibco) and cortices were dissected in ice cold HBSS. After the removal of debris (130-107-677, Miltenyi Biotec), cells were resuspended with 1 mL “re-expression medium” containing NeuroBrew-21 (1:50 in MACS neuro Medium) (130-093-566 and 130-093-570, Milteny Biotec) and 200 mM L-glutamine (1:100, G7513, Sigma) at 37 °C for 30 min. Cells were then incubated with Fc-receptor blocker for 10 min at 4 °C (provided with CD140 microbeads kit), followed by a 15 min incubation with 10 µL microbeads mixture composed of CD140 (130-101-502), NG2 (130-097-170) and O4 (130-096-670) in 1:1:1 at 4 °C. For the purity assessment, cells were re-suspended with 1 mL “re-expression medium,” seeded on the coverslips which were coated with polyonithin (P2533, Sigma). After 2 hrs incubation at 37 °C with 5 % CO_2_, cells were processed for immunocytochemistry against PDGFRα and DAPI. For qRT-PCR or Western blot analysis, MACS sorted OPCs were lysed by RIPA buffer (89900, Thermo Scientific).

#### Western blot analysis

Deeply anesthetized mice were perfused with cold PBS. Medial prefrontal cortices (mPFC) and corpus callosa (cc) were dissected in ice cold PBS. Tissues were homogenized with sucrose lysis buffer (320 mM sucrose, 10 mM Tris-HCl, 1 mM NaHCO_3_ (pH=7.4); 1 mM MgCl_2_), and MACS-orted OPCs were lysed with RIPA buffer. Both buffers were supplemented with 1 X protease inhibitors (05892970001 Roche) and 1 X phosphatase inhibitor (04906837001, Roche). Protein (5 µg) was blotted onto nitrocellulose transfer membranes (QP0907015, qpore). Membrane was blocked with 5 % non-fat milk or 5 % BSA (A7906, Sigma) diluted in 0.1 % TBST. Primary antibodies were diluted with corresponding blocking buffer. Primary antibodies used for Western blot are listed in the **Supplementary table 3.**

Secondary antibodies were: HRP anti-mouse (1:2000,) and anti-rabbit (1:2000). Membranes were illuminated with WesternBright Chemilumineszenz Substrat Quantum kit (541015, Biozym) and documented with ChemiDoc-MP.

#### Quantitative real time PCR

Brain tissue or the MACS-sorted OPCs were homogenized as described above. NucleoSpin RNA Plus XS kit (740990.50, Macherey-Nagel) was used to extract mRNA and Omniscript kit (205113, QIAGEN) was used for reverse transcription. RT-PCR was performed using EvaGreen (27490, Axon) kit. Primer sequences for qRT-PCR are listed in **Supplementary table 4**.

#### Electrophysiology

##### Slice preparation

Mice were anesthetized by isoflurane before decapitation, and brain was quickly isolated and immersed in an ice-cold, oxygenated (5 % CO_2_/95 % O_2_, pH=7.4) solution containing (in mM) 87 NaCl, 3 KCl, 25 NaHCO_3_, 1.25 NaH_2_PO_4_, 3 MgCl_2_, 0.5 CaCl_2_, 75 sucrose and 25 glucose. Coronal or semi-sagittal slices in 300 µm thickness were prepared with vibratome (Leica VT 1200S, Nussloch, Germany) and transferred to a nylon basket slice holder for incubation in artificial cerebral spinal fluid (ACSF) containing (in mM) 126 NaCl, 3 KCl, 25 NaHCO_3_, 15 glucose, 1.2 NaH_2_PO_4_, 2 CaCl_2_, and 2 MgCl_2_ at 32 °C for 0.5 h. Subsequently, slices were removed from the water bath and kept at RT with continuous oxygenation prior to use.

##### IPSCs and EPSCs recordings

Semi-sagittal slices were transferred to the recording chamber that was continuously perfused with oxygenated ACSF containing 1 MgCl_2_ and 2.5 CaCl_2_ at a flow rate of 2-5 mL/min. During recordings, 50 μM strychnine and 50 μM picrotoxin were added to block inhibitory synaptic transmission. Pyramidal neurons were identified morphologically (Axioskop 2 FS mot, Zeiss, Jena, Germany) with a 40x water immersion objective. Tissue was detected with a QuantEM 512SC camera (Photometrics, Tucson, USA). Whole-cell membrane currents were recorded by an EPC 10 USB amplifier (HEKA, Lambrecht, Germany), low pass filtered at 3 kHz and data acquisition was controlled by Patchmaster software (HEKA). The resistance of patch pipettes (7–9 ΩM) were fabricated from borosilicate capillaries (OD: 1.5 mm; Sutter, USA) using a Micropipette Puller (Model P-97, Sutter Instrument Co., CA). Patch pipettes were filled with the solution containing (in mM) 125 cesium gluconate, 20 tetraethylammonium (TEA), 2 MgCl_2_, 0.5 CaCl_2_, 1 EGTA, 10 HEPES and 5 Na_2_ATP (pH=7.2). Spontaneous excitatory and inhibitory postsynaptic currents (sEPSCs, sIPSCs) of pyramidal neurons in mPFC were recorded for 40 s in voltage-gated mode at a holding potential of −70 mV and +30 mV, respectively. Currents above 10pA were analyzed with MATLAB. EPSCs of pyramidal neurons in the layer V of mPFC were recorded by stimulating presynaptic axons in the layer II/III of mPFC with a concentric bipolar microelectrode (MicroProbes, USA). Stimulus duration was 200 µs. To estimate short-term plasticity, 40 trains of stimulation were applied to induce synaptic depression, and the depressing trains were repeated with frequency of 10 Hz and an interval of 20 s to allow for recovery from synaptic depression. The stimulation threshold was applied from 30 µA to 80 µA. IPSCs were conducted in the same region, however with a holding potential of 0 mV.

##### Compound action potential recordings

Compound action potentials were recorded in the corpus callosum as previously described (Crawford et al., 2009). Micropipettes with 1-3 MΩ resistance were filled with ACSF. Inward responses were evoked in current-clamp mode by varying the intensity of stimulus pulses (0.2-4.0 mA) at 1 mm distance between recording and stimulation electrodes, with a stimulus duration of 200 μs. The sample sweeps were acquired every 5 s. Conduction velocity was estimated by changing the distance from 2.5 mm to 0.5 mm between the stimulating and recording electrodes with a constant stimulus. To enhance the signal to noise ratio, we averaged at least 15 successive sweeps. Data analysis was performed with Igor pro 6.3.7.2 (WaveMetrics, Oregen, US).

All the experiments were conducted at RT (22 – 24 °C).

##### Data analysis

Data generated by Patch Master were loaded into MATLAB (Mathworks, MA, USA) with a module adapted from sigTOOL (Lidierth, 2009). Evoked EPSC and IPSC traces from the same cells were manually checked and pooled. The average of EPSC/IPSC traces from each cell was used for analysis. Data analysis was performed using routines that were custom written in Matlab.

#### EEG telemetry and analysis

Telemetric EEG transmitter implantation was adapted from Bedner and colleagues ^7^. Mice were implanted with telemetric EEG transmitters (DSI PhysioTel® ETA-F10, Harvard Biosciences, Inc. Holliston, Massachusetts, USA) between 8 and 10 weeks of age. The animals were placed in a stereotaxic frame (Robot stereotaxic, Neurostar, Tübingen, Germany) for implantation of depth electrodes at 3.4 mm posterior to bregma and bilaterally 1.6 mm from the sagittal suture. After post-surgical care and recovery, cages were placed on individual radio receiving plates (DSI PhysioTel® RPC-1, Data Sciences International, St. Paul, USA) for synchronized EEG and video recording (MediaRecorder Software, Noldus Information Technology, Wageningen, Netherlands). EEG and video recording were performed for 24 hrs.

##### Data analysis

EEG traces were analysed with the Neuroscore software (Version 3.3.1., Data Sciences International, St. Paul, USA). After applying a general Notch filter (50 Hz band stop), relative power band values were determined with order 13 fast-Fourier transform in 10s epochs and later averaged per hour of recording. Statistical analysis was performed with GraphPad Prism.

#### Behavioral analysis

Several behavioral tests were performed with the same cohorts of mice (male, 9-week-old). The order of the tests was designed from low to high invasiveness to reduce the interference from the prior tests. The chamber or tested square was wiped with 75 % ethanol before tests to remove odors.

##### Nest building test

The nest building test was carried out as previously described ^8^. Mice were moved into new single cages in the behavior testing room for habituation. After 24 h, a piece of pressed cotton in size of 7×5 cm2 was placed in the cage. After 14 h, the shape and weight of the cotton was recorded and unbiasedly scored according to the criteria ^8^.

##### Open field test

The mice were put in the open field maze, which measured 50 cm (length) x 50 cm (width) x 38 cm (height). Mice could move and explore freely for 10 min in the open field square. In each test, the single mouse was put in the center of the square arena. The videos of these 10 min were recorded (USB webcam) and analyzed (EthoVision XT 11.5, Noldus Technology). Duration time in the center area (s), moved distance (cm) and speed (cm/s) were determined ^9^

##### New object recognition test

The mice could freely explore the two identical objects placed at a distance of 8.5 cm from the side walls in two opposite corners of the apparatus for 10 min. After 40 min, one of the objects was replaced by a novel one, and the test mouse was allowed to explore again. Preference index was defined as the percentage of the time exploring one identical object within the total time exploring both objects. Recognition index was defined as the percentage of the time exploring the novel object among the total time of exploring both objects.

##### Three chamber social behavior test

The three-chamber box was employed for social behavioral studies ^10^. Three chambers were equally sized and separated by two walls evenly distributed in the box. A square door at the bottom center of each door allowed free running of the mice within the three chambers. Two empty wire cages were placed at the side chambers, leaving the center chamber empty. For habituation, mice were kept in the center chamber for 10 min, followed by a 10 min-habituation session with access to all three chambers. For the sociability test, the test mouse was placed in the center chamber, while a mouse of similar age was kept under the stainless wire cage in one of the side chambers. The other chamber contained an empty wire cage. For 10 min, the test mouse could select freely all three chambers. For the social novelty test, a novel mouse was placed under the other empty cage being an unfamiliar mouse, and the prior mouse as familiar one. The test mouse was again allowed to freely explore both animals for 10 min. The experiment was recorded with video camera and the time that the test mouse spent in each chamber and the time of sniffing was analyzed.

#### Cell culture

##### Cell line Oli-neu

The murine oligodendroglial precursor cell line Oli-neu ^11^ was kindly provided by Professor Jacqueline Trotter (University of Mainz). Undifferentiated Oli-neu cells were incubated at 37 °C and 5 % CO_2_ in poly-L-lysine (Merck) coated cell culture flasks (Greiner Bio one) for expansion or cell culture dishes (Greiner Bio one) for experiments. Sato medium consisting of DMEM high glucose medium (Fisher Scientific) with supplementation of 10 µg/ml transferrin (Sigma), 10 µg/ml insulin (Santa Cruz), 100 µM putrescine (Sigma), 200 nM progesterone (Sigma), 500 nM tri-iodo-thyrodine (Sigma), 220 nM sodium selenite (Sigma), 520 nM L-thyroxine (Sigma) and 1.5 % normal horse serum (Fisher Scientific) was used for culturing and proliferation of cells. For experiments, 2×10^5^ cells were seeded in a 60 mm Petri dish. 48 hours later, the medium was changed to fresh, with or without CGP 55845 (50 µM) for 24 hours. Cells were lysed with Qiazol (Qiagen) for qRT-PCR in a blind manner. Conditional medium of control cells was collected for further neuronal treatment.

##### Primary culture of cortical neurons

Cortical neurons were isolated from p0 pups (C57BL/6 mouse strain). Briefly, the cortex was dissected from the whole brain in ice cold Earle’s Balanced Salt Solution (EBSS, Gibco). The cortex was then digested with 35 units papain (Worthington,NJ) for 45 min at 37 °C, followed by gentle mechanical trituration. 1×105 cells were seeded on 25 mm glass coverslips in 24-well culture plates for further immunostaining. The glass coverslips were pre-coated with a mixture of coating solution containing 17 mM acetic acid, poly-D-Lysine (Sigma, P6407) and collagen I (Gibco, A1048301). Neurons were cultured in NBA culture medium that contained 10 % FCS, 1 % penicillin-streptomycin, 1 % GlutaMAX and 2 % B-27 supplement (Gibco) for 7 d at 37 °C with 5 % CO_2_ incubator before the experiment. To examine the effect of TWEAK on neurons, conditioned Oli-neu culture medium supernatant or Sato, was applied to neuron NBA culture medium, with a ratio of 1:2 to NBA resulting in 1 ml medium per 24-well and 2 ml medium per 6-well. Cells were collected after 6 h with different culture condition with or without 20 µM TWEAKR inhibitor L524-0366 (509374, Calbiochem) for further analysis.

#### Image acquisition and analysis

Brain slices were scanned with the fully automated slide scanner AxioScan.Z1 (Zeiss, Jena) and LSM 710 confocal microscope (Zeiss, Jena). Cell counting was manually performed without bias by using ZEN software (Zeiss, Jena). The lengths of nodes and paranodes were measured using the ‘straight’ tool of Fiji software. The Imaris (version 9.6) with functions of volume analysis and vesicle classification were used for MBP-PV (Fig. 1M, N) and vGAT study (Fig. F-I). Cell counting and Imaris analysis were carried out in a blind manner.

#### Bromodeoxyuridine assay

Adult animals received 1 mg/ml bromodeoxyuridine (BrdU) (B5002, Sigma-Aldrich, St. Louis, MO) dissolved in drinking water for seven consecutive days. For juvenile mice, single shot of 10 mg/ml BrdU dissolved in 0.9 % NaCl were intraperitoneally injected to the p13 pups and analyzed at p14.

#### Electron microscopy

##### Perfusion & dissection

Anesthetized (Ketamine/Xylazine in 0.9% NaCl solution (all: Bayer, Leverkusen, Germany)) mice were intracardially perfused with buffered heparin (B. Braun, Melsungen, Germany) solution (end concentration 50 IU/mL) followed by perfusion fixation with 4% (w/v) paraformaldehyde and 0.5% (w/v) glutaraldehyde in 0.1M cacodylate buffer. During perfusion, the solutions were gradually cooled down from 37°C to ice cold temperatures. Whole brains were dissected, immersed in fixative solution for 24h, and stored in 0.1M cacodylate buffer at 4°C until further use.

##### Slicing

For vibratome slicing, whole brains were embedded in 10% gelatin and immersed in ice cold 0.1M cacodylate buffer. 500 µm frontal sections were made using a vibratome (VT1000S, Leica Microsystems, Wetzlar, Germany) at a frequency of 80Hz and 0.1mm/s speed. Slices containing the region of interest were determined using a mouse brain atlas ^12^, and particular brain areas (i.a. corpus callosum) were meticulously cut out using a fine scalpel.

##### Embedding & examination

The specimen of the different brain regions were embedded for electron microscopy (1975) ^13^. In short, samples were repeatedly rinsed in 0.1M cacodylate buffer, osmicated in 2% osmium tetroxide in 0.1M cacodylate buffer for 1h, washed in distilled water, and dehydrated in an ascending series of ethanol (70% to 100%) and acetone (100%, water-free). An embedding in Epon resin (EMS, Hatfield, PA, USA) followed. After thorough polymerization, semi-in (300nm) and ultra-thin (65nm) sections were cut using an ultra-microtome (EM UC7, Leica Microsystems, Wetzlar, Germany). Semi-thin sections were stained with Richardson blue staining solution, and examined under a light microscope (DM2500, Leica Microsystems, Wetzlar, Germany). Ultra-thin sections were contrasted with 5% lead citrate for 5min, examined with a transmission electron microscope (Tecnai G2, FEI, Hillsboro, OR, USA), and randomly selected areas were documented with a digital camera (MegaView III, Olympus, Shinjuku, Japan).

#### Statistics

Prior to statistical analysis, data were tested for outlier identification and Gaussian distribution with GraphPad Prism 8. Statistical differences were indicated in the figures and legends. Each data point in the graphs indicates biological individuals. Exceptions are indicated in the figure legends.Data are shown as mean±SEM, except for violin plots (Fig. 2i and Fig. 4b). The median and quartiles are indicated as thick and thin dashed lines, respectively.

---

**Supplementary table 1:** Primary antibodies used for immunostaining.

| **Antibodies** | **Host** | **Dilutions** | **Cat.No.** | **Company** | **PRID** |
| --- | --- | --- | --- | --- | --- |
| **PDGFRα** | goat | 1:500 | AF1062 | R&D Systems | AB_2236897 |
| **adenomatous polyposis coli clone 1(CC1)** | mouse | 1:200 | OP80 | Calbiochem | AB_2057371 |
| **Olig2** | rabbit | 1:500 | AB9610 | Millipore | AB_570666 |
| **MBP** | mouse | 1:500 | SMI99 | Biolegend | AB_10120130 |
| **Parvalbumin** | rabbit | 1:1000 | PV 25 | Swant | AB_10000344 |
| **Parvalbumin** | mouse | 1:500 | P3088 | Sigma | AB_477329 |
| **Caspr** | rabbit | 1:500 | ab34151 | abcam | AB_869934 |
| **DsRed** | rabbit | 1:1000 | 632496 | Clontec | AB_10013483 |
| **BrdU** | rat | 1:1000 | ab6326 | abcam | AB_305426 |
| **Cleaved Caspase-3** | rabbit | 1:200 | 9661 | Cell Signaling Technology | AB_2341188 |
| **Cleaved Caspase-3** | mouse | 1:100 | STJ9744 | St. John's Laboratory |  |
| **GAD67** | mouse | 1:500 | MAB5406 | Millipore | AB_2278725 |
| **NeuN** | mouse | 1:500 | MAB377 | Millipore | AB_2298772 |
| **CTIP2** | rat | 1:200 | 650601 | Biolegend | AB_10896795 |
| **TBR1** | rabbit | 1:500, | 49661 | Cell Signaling Technology | AB_2799364 |
| **vGAT** | mouse | 1:500 | 13002 | Synaptic Systems | AB_887871 |
| **FN14** | mouse | 1:50 | 314108 | Biolegend | AB_2810477 |

**Supplementary table 2:** Secondary antibodies used for immunostaining.

| **Antibodies** | **Fluorophore** | **Cat. No.** | **Company** |
| --- | --- | --- | --- |
| **Donkey anti-mouse** | Alexa Fluor® 488 |  | Thermofischer |
|  | Alexa Fluor® 546 | A10036 |  |
|  | Alexa Fluor® 647 | A31571 |  |
|  | DyLight® 755 | SA5-10171 | Invitrogen |
| **Donkey anti-rabbit** | Alexa Fluor® 488 | A21206 | Thermofischer |
|  | Alexa Fluor® 546 | A10040 |  |
|  | Alexa Fluor® 647 | A31573 |  |
|  | Alexa Fluor® 790 | A11374 |  |
| **Donkey anti-goat** | Alexa Fluor® 488 | A11055 | Thermofischer |
|  | Alexa Fluor® 546 | A11056 |  |
|  | Alexa Fluor® 647 | A21447 |  |
|  | Alexa Fluor® 750 | ab175744 | abcam |
| **Donkey anti-rat** | DyLight-755 | SA5-10031 | Thermofischer |
| **DAPI (25ng/ml)** |  | A10010010 | biochmica |

Secondary antibodies were diluted 1:1000 with the blocking buffer.

**Supplementary Table 3.** Primary antibodies used for Western blot.

| Antibodies | Host | Dilutions | Cat.No. | Company | PRID |
| --- | --- | --- | --- | --- | --- |
| GABA<sub>B1</sub>R | mouse | 1:500 | ab55051 | abcam | AB_941703 |
| GAPDH | mouse | 1:2000 | G8795 | Sigma | AB_1078991 |
| MBP | mouse | 1:1000 | SMI99 | Biolegend | AB_10120130 |
| Tubulin-α | rabbit | 1:2000 | T6074 | Sigma | AB_477582 |
| TWEAK | rabbit | 1:500 | NBP1-76695 | Novus Biologicals | AB_11036308 |

**Supplementary Table 4.**
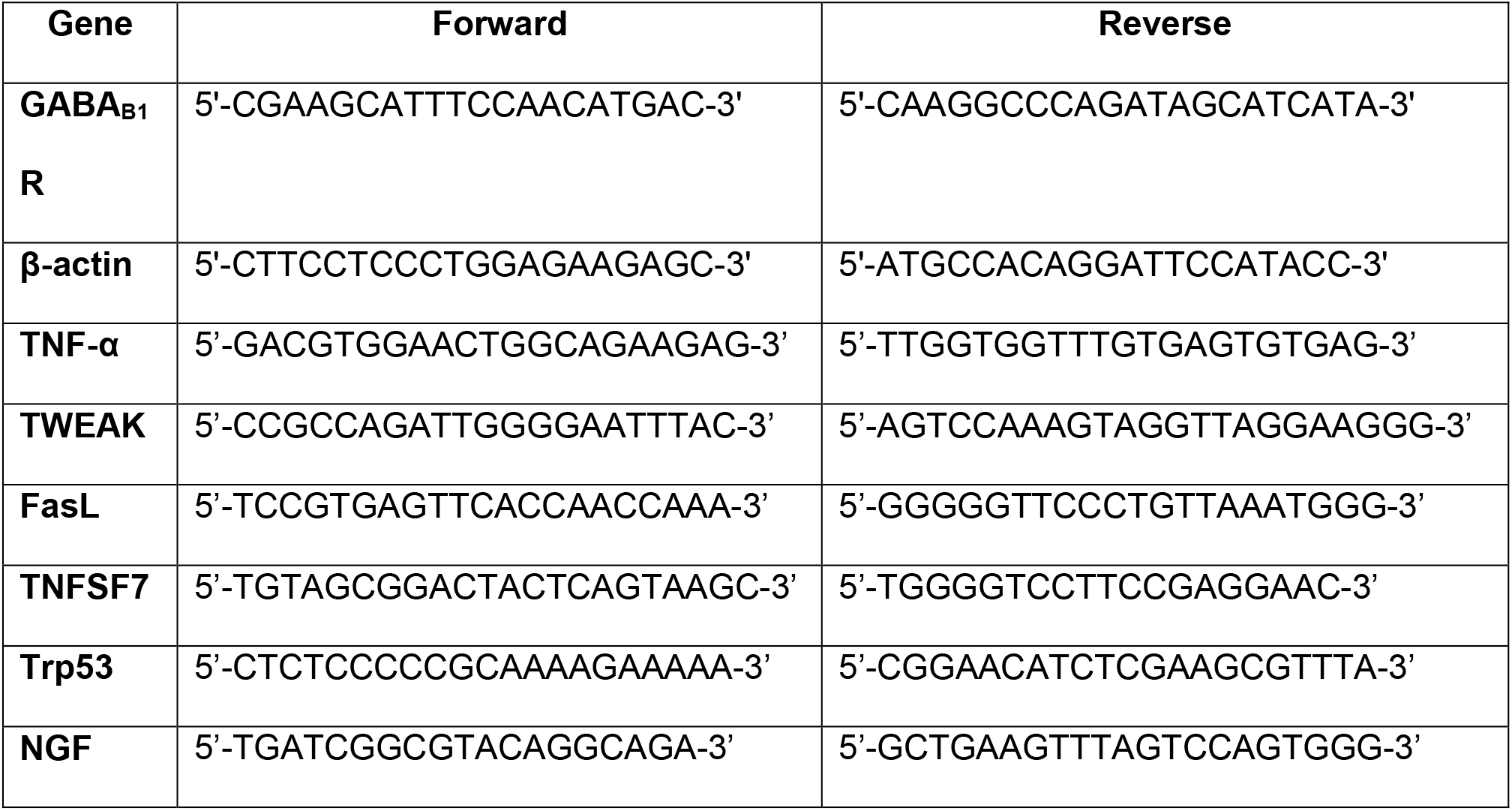
Primers used for qRT-PCR.

